# Ultra-fast and scalable high-resolution full-length single-cell RNA sequencing using CHART-seq

**DOI:** 10.64898/2026.09.18.752546

**Authors:** Wenyi Zhang, Aiqun Chen, Kaiqiang Ye, Handong Wang, Jiahao Shen, Zhe Jiao, Yunxia Guo, Yiming Xu, Dongmei Zhang, Yan Huang, Liyong He, Xiaofei Gao, Xiangwei Zhao

## Abstract

Plate-based full-length single-cell RNA sequencing resolves transcript structure details but remains difficult to scale because each cell usually requires a separate library. Here we developed CHART-seq (**C**ombinatorial **H**eteroduplex **A**ssay via **R**ecombinant **T**n5), which uses orthogonally indexed Tn5 complexes to tagment RNA/cDNA heteroduplexes and permits early sample pooling. The workflow processed up to 96 cells per library, was compatible with 384-well expansion, and completed library preparation within 3 h at a reagent cost below US<$>1 per cell. At matched sequencing depth, CHART-seq detected more genes and annotated isoforms than Smart-seq2, Smart-seq3, Flash-seq and SHERRY2, while retaining broad genebody coverage and reproducible expression estimates. In the CHART-seq results of vascular smooth muscle cells, TGF-β1 pretreatment before PDGF-BB exposure partly restored contractile features, suppressed a PDGF-associated inflammatory programme, and induced a distinct metabolic–matrix response with coordinated transcript-usage changes. These biological findings remain exploratory because independent biological replicates were unavailable. CHART-seq provides a rapid, scalable route to full-length single-cell transcript profiling with gene-programme and candidate isoform resolution.

**Graphical Abstract:** 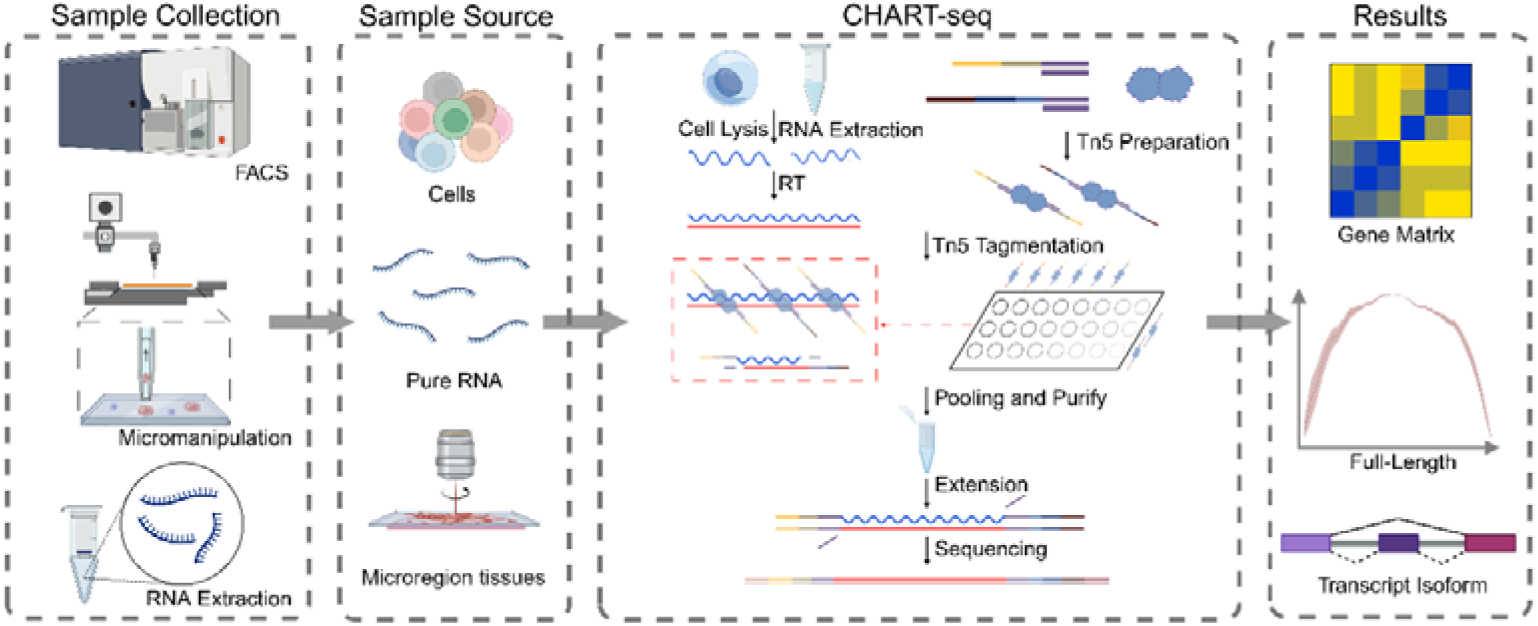

## Introduction

Technical choices in single-cell RNA sequencing have long been constrained by the trade-off between cell throughput and the amount of transcript information recoverable per cell(1–4). Droplet-based 3′ or 5′ end counting enables gene expression profiling across large numbers of cells. Still, it struggles to simultaneously retain intra-transcript sequence information, thereby limiting the resolution of transcript isoforms, allele-specific expression, and transcriptional variation(5,6). Plate-based full-length single-cell RNA sequencing exchanges lower cell throughput and a heavier library-construction burden for higher sensitivity and more complete gene-body coverage, and therefore remains an important technical route for dissecting single-cell transcript structure(7–9).

However, traditional methods’ amplification and library-preparation workflow is time-consuming and difficult to scale(10). Smart-seq3 introduced unique molecular identifiers (UMIs) to improve molecule counting, and Flash-seq shortened the workflow through redesigned reverse-transcription chemistry(11–13). SHERRY and SHERRY2 further simplified library construction by applying Tn5 transposase directly to RNA–DNA heteroduplexes(14,15). Despite these advances, most plate-based workflows still preserve a one-cell–one-library architecture until late in the protocol. Consequently, processing cost and library-to-library variation increase with cell number. However, these improvements primarily increase the efficiency of the per-well reaction; during downstream amplification and library processing, samples still need to remain physically separated well by well for a considerable period(16,17). As the number of cells increases, the number of handling steps, reagent consumption, and batch effects inevitably increase in parallel(18,19). This raises a key question: whether sample identity can be encoded at the stage of tagging RNA/cDNA hybrids, allowing samples to be pooled at an early step while preserving low-input sensitivity and full-transcript information(20,21).

We reasoned that orthogonally barcoded Tn5 complexes could identify individual wells during heteroduplex tagmentation, thereby permitting early pooling. This design would retain the advantages of plate-based full-length profiling while reducing the number of purification and amplification reactions. We therefore developed CHART-seq, a combinatorial heteroduplex-tagmentation workflow in which two indexed Tn5 complexes introduce well-specific barcode combinations before products are pooled, purified, gap-filled and amplified as a shared library.

In this article, we first established the chemistry using purified HepG2 total RNA, optimised the workflow at 1 ng and 10 pg input, and benchmarked its analytical performance against four established full-length methods at matched read depth.

Meanwhile, coronary artery disease is one of the most prevalent cardiovascular diseases worldwide and poses a serious threat to human health. Percutaneous coronary intervention (PCI) is a core therapeutic approach for coronary artery disease and has been widely used in clinical practice. However, in-stent restenosis, the most common cause of PCI failure, remains a major challenge for both patients and clinicians(22–24). Based on clinical observations and clinical research data, we found that patients with diabetes mellitus are more prone to in-stent restenosis after PCI(25,26). Previous studies have reported that the classic mechanism underlying in-stent restenosis is the phenotypic switching of vascular smooth muscle cells (VSMCs) from a contractile phenotype to a synthetic phenotype, with PDGF-BB released after endothelial injury as the classic inducing factor(27,28). In the diabetic environment, however, VSMCs tend to undergo a more fibroblast-like VSMC transition, for which the classic inducing factors are PDGF-BB combined with TGF-β1(29,30). To investigate the differences among contractile, synthetic, and fibroblast-like VSMC phenotypes, we used CHART-seq for exploration.

## Results

### CHART-seq

The core design of CHART-seq is to shift sample-identity labelling from downstream PCR amplification upstream to the RNA/cDNA hybrid tagging step. Poly(A)+ RNA from lysed cells or purified RNA is first reverse-transcribed using oligo(dT) primers; subsequently, two Tn5 transposomes carrying Read 1 and Read 2 adaptors, respectively, together with orthogonal well barcodes, are used to tag the RNA/cDNA hybrids within each well. After in-well encoding is completed, all reactions can be pooled, and column purification, Bst 3.0 extension, and PCR amplification are then carried out in a single tube (Fig. 1A). This workflow shortens the library-construction time from purified RNA to approximately 2.5 h, and keeps the entire procedure for cell samples within 3 h. The absence of preamplification after RT reduced the opportunity for early PCR distortion, while the UMI supported computational removal of amplification duplicates. Because the barcode combination was introduced before pooling, up to 96 samples could be processed in one library in the tested design, with a direct path to 384-well scaling through additional index combinations.

**Figure 1:**
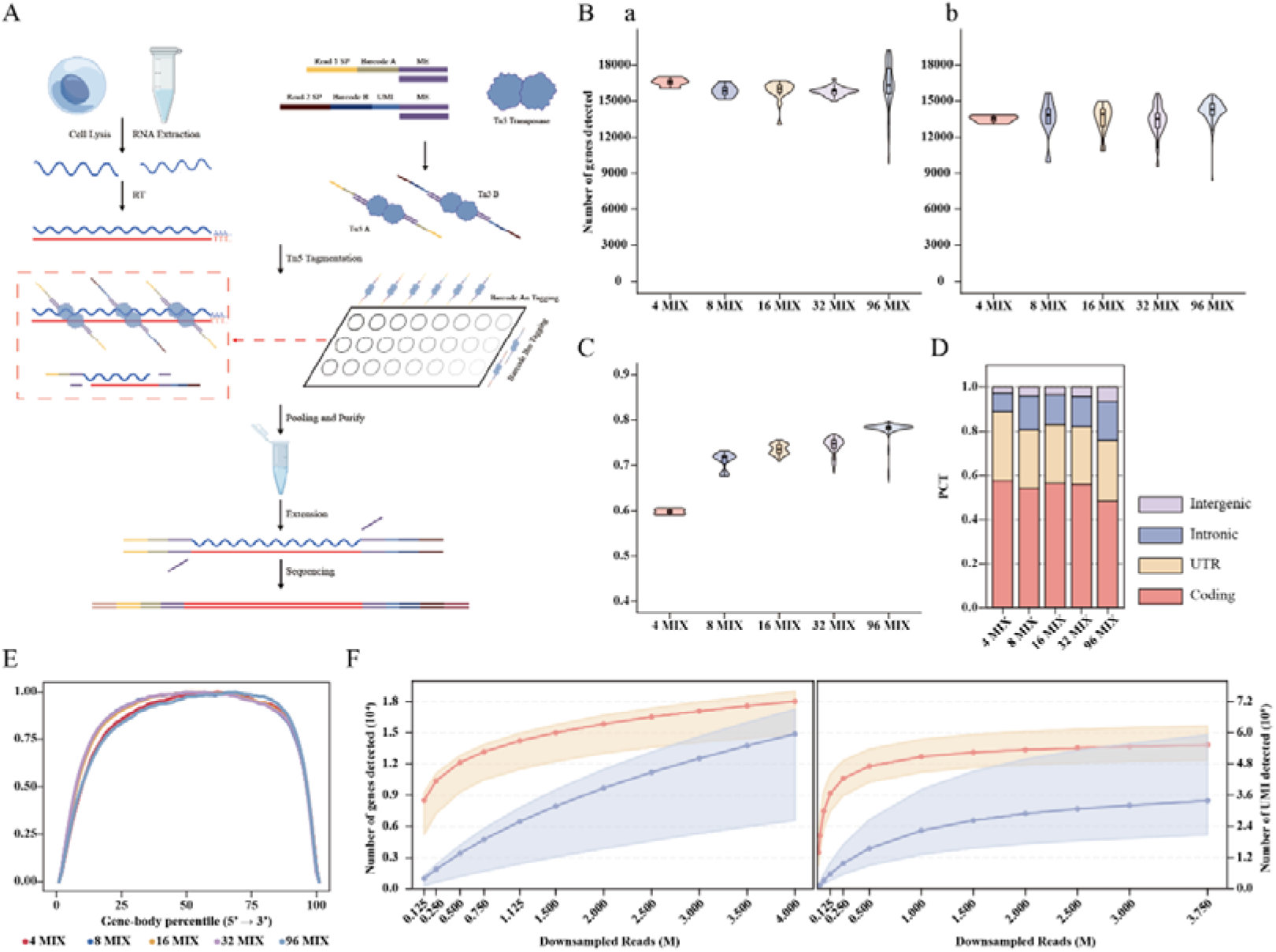
Overview of CHART-seq. **A**, Five-step CHART-seq workflow. **B**, Genes detected from 1 ng (**a**) and 10 pg (**b**) HepG2 total RNA in 4-, 8-, 16-, 32-, and 96-sample pools; samples were downsampled to mean depths of 2 million and 500,000 reads, respectively. **C**, Pairwise cell-to-cell correlation for 10 pg RNA across pooling formats. **D**, Genomic-region distribution of CHART-seq reads from 10 pg RNA. **E**, Gene-body coverage for 10 pg RNA. **F**, Gene-and UMI-saturation curves for 1 ng (left) and 10 pg (right) RNA; ribbons show the observed sample range.

We initially used 1 ng of HepG2 total RNA to establish the order and operating conditions of purification, extension, reverse transcription, heteroduplex tagmentation and library amplification (Supplementary Figure S1). However, when the input amount is reduced to 10 pg, reaction efficiency and purification recovery become prominent factors limiting library complexity. To overcome these problems, we optimised reverse transcription, tagmentation and purification at an input of 10 pg total RNA, approximating the RNA content of a mammalian cell (Supplementary Figures S2–S4). Purification was particularly limiting because polyadenylated RNA represents only a small fraction of total RNA and low-input heteroduplex products are readily lost on solid-phase matrices. Adding a 100-bp synthetic double-stranded support DNA improved gene and UMI recovery (Supplementary Figure S4A, B). The sequence was derived from Escherichia coli and selected to lack detectable similarity to the human and mouse reference genomes, thereby providing carrier mass without intentionally contributing alignable mammalian reads.

Early pooling of samples may lead to barcode imbalance, dilution of low-input samples, or reduced library complexity. To examine this risk, equal amounts of 1 ng or 10 pg HepG2 total RNA were distributed across 4, 8, 16, 32, and 96 barcode combinations, and gene detection was compared at a fixed per-sample sequencing depth (Fig. 1B). At 1 ng input and a mean depth of 2 million reads per sample, more than 15,000 genes were detected. At 10 pg input and 500,000 reads per sample, more than 11,000–12,000 genes were detected. Pairwise Kendall correlations among equal-input 10 pg samples averaged above 0.75 across pooling formats (Figure 1C), indicating that the technique maintains stable and robust reproducibility as the pooling scale increases. More than half of aligned bases mapped to protein-coding regions and more than 80% mapped to exons, whereas intergenic reads remained below 5% (Figure 1D). This indicates that the majority of valid reads are derived from annotated transcript regions. Gene-body coverage was broad, with a coverage-uniformity value above 0.8, supporting the acquisition of sequence information spanning the transcript body (Figure 1E).

Sequencing-saturation analysis separated the effects of input mass and read depth (Figure 1F). At 1 ng input, approximately 15,000 genes and 350,000 UMI-resolved fragments were recovered at 2 million reads, increasing to more than 18,000 genes and 550,000 UMIs at 4 million reads. At 10 pg input, approximately 11,000 genes and 150,000 UMIs were recovered at 500,000 reads; gene discovery approached saturation above 15,000 genes by 2 million reads, while UMI recovery continued to increase beyond 250,000 molecules. These data indicate that relatively shallow sequencing is sufficient for broad gene detection at single-cell-equivalent input, whereas deeper sequencing mainly improves molecule recovery and transcript-level resolution.

### Benchmarking against other full-length single-cell RNA-sequencing methods

We compared CHART-seq in HEK293T cells with Smart-seq2, Smart-seq3, Flash-seq and SHERRY2 after harmonising the computational workflow and downsampling each cell to 500,000 reads (Figure 2). CHART-seq is a full-length transcriptome sequencing method that completes the entire procedure, from RNA denaturation to finished library construction, within 3 h (Figure 2A). It detected a mean of 10,287 genes and 21,920 annotated transcript isoforms per cell, the highest values among the five datasets under the matched-depth analysis (Figure 2B; Supplementary Figure S5A). CHART-seq also recovered longer transcripts and a higher fraction of transcripts exceeding 2 kb and 5 kb than several comparator datasets (Supplementary Figure S5B–F), consistent with broad gene-body representation.

**Figure 2:**
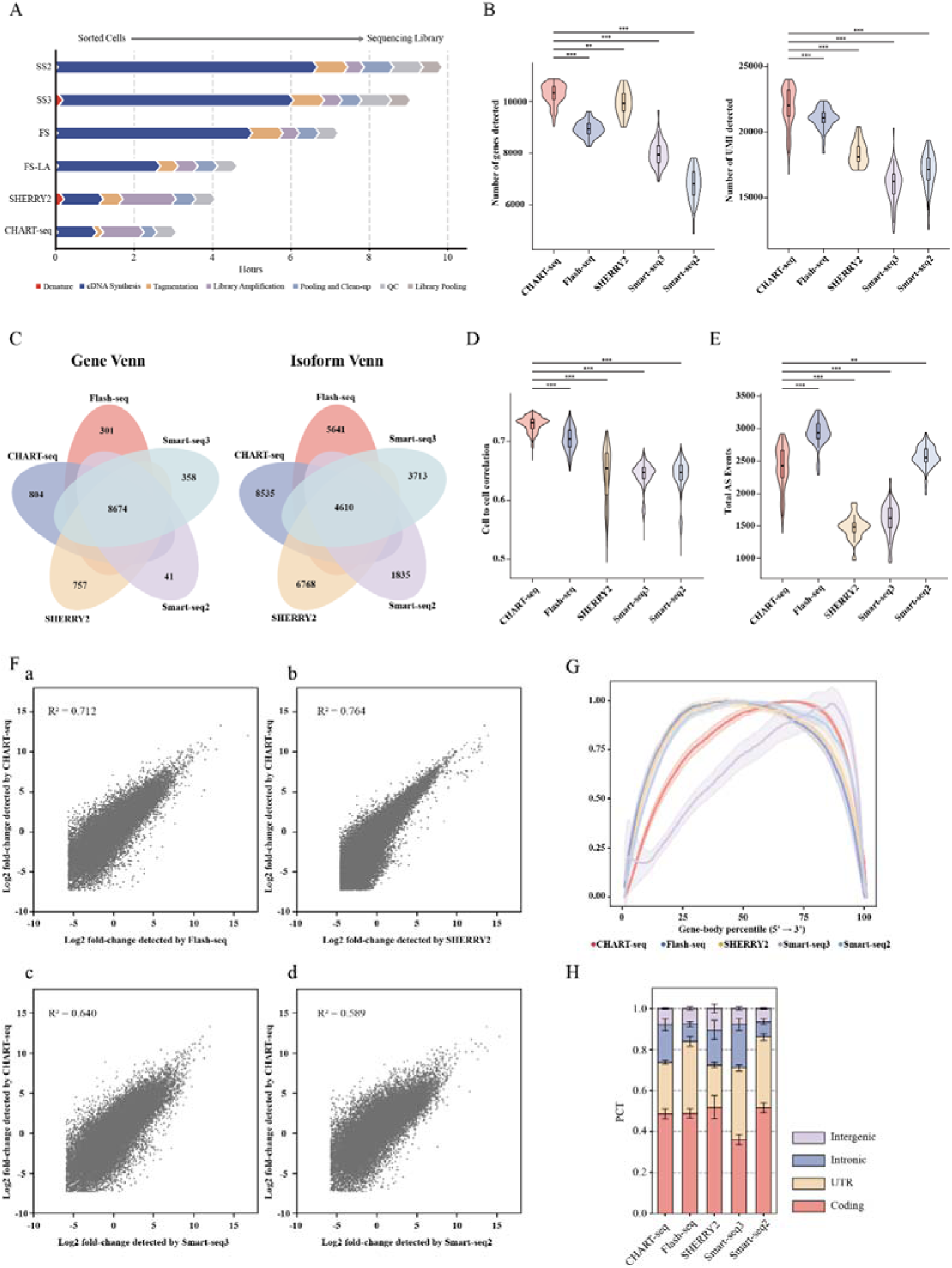
Benchmark with other Full-length single-cell RNA-seq methods in HEK 293T cells. All cells were downsampled to 500,000 reads, CHART-seq contains 160 cells (8 libraries), Flash-seq, Smart-seq2 and Smart-seq3 contain 50 cells, and SHERRY2 contains 24 cells . A, Time from RNA denaturation to a sequencing-ready library for CHART-seq, Smart-seq2, Smart-seq3, Flash-seq, and SHERRY2. B, Detected genes (left) and annotated transcript isoforms (right). C, Five-method overlap of detected genes (left) and isoforms (right). D, Within-method cell-to-cell expression correlation. E, Total detected alternative-splicing events. F, Concordance of gene-expression fold changes between CHART-seq and each comparator among genes detected by both methods. G, Gene-body coverage. H, Fractions of reads assigned to coding, untranslated, intronic and intergenic regions.

The five methods shared 8,674 detected genes, but only 4,610 annotated isoforms (Figure 2C). Thus, gene-level detection was substantially more concordant than transcript reconstruction. This divergence is expected for short-read sequencing, in which transcript identities are inferred from fragments rather than observed as intact molecules. Isoform counts should therefore be interpreted as annotation-supported estimates, not as direct single-molecule validation(31–34).

CHART-seq also showed high cell-to-cell expression concordance (Figure 2D). The 160 CHART-seq cells were generated in eight pooled libraries, whereas comparator protocols generally retained individual libraries for each cell. Early pooling may reduce variation introduced by repeated purification and amplification reactions, although the cross-study design does not isolate chemistry from differences in sample handling, operator, sequencing run or data source. CHART-seq detected fewer alternative-splicing events than Flash-seq, a similar number to Smart-seq2, and more than Smart-seq3 and SHERRY2 (Figure 2E; Supplementary Figure S6). Gene-expression fold changes showed the greatest concordance between CHART-seq and SHERRY2 (R²=0.764; Figure 2F), consistent with the shared principle of direct RNA–DNA heteroduplex tagmentation.

CHART-seq retained broad gene-body coverage but showed a modest 3′ bias compared with the best-performing comparator (Figure 2G). A plausible contributor is partial RNA degradation during DNase I treatment after cell lysis, despite the presence of RNase inhibitor. Genomic-region analysis further showed that CHART-seq and SHERRY2 had coding-region fractions similar to other methods but higher intronic fractions than Smart-seq2 and Flash-seq, without a corresponding increase in intergenic reads (Figure 2H). This pattern is compatible with greater recovery of unspliced or nascent RNA, although dedicated metabolic-labelling or nuclear-fractionation experiments would be required to establish that interpretation(8,33). Under the tested pooling design, CHART-seq reduced reagent cost to below US<$>1 per cell and allowed hundreds of cells to be processed within one working session (Supplementary Figure S7).

### Combined stimulation partly restores PDGF-BB-induced VSMC states

We profiled untreated VSMCs (G1), cells stimulated with PDGF-BB (G2), and cells pretreated with TGF-β1 before PDGF-BB exposure (G3; Figure 3A, B). The design comprised 40 intended cells per group distributed across three technical libraries. After correction of six surplus barcode records in the second and third libraries and blinded removal of cells with extreme complexity, the primary analysis retained 99 cells (G1, 27; G2, 37; G3, 35; Supplementary Figure S8).

**Figure 3:**
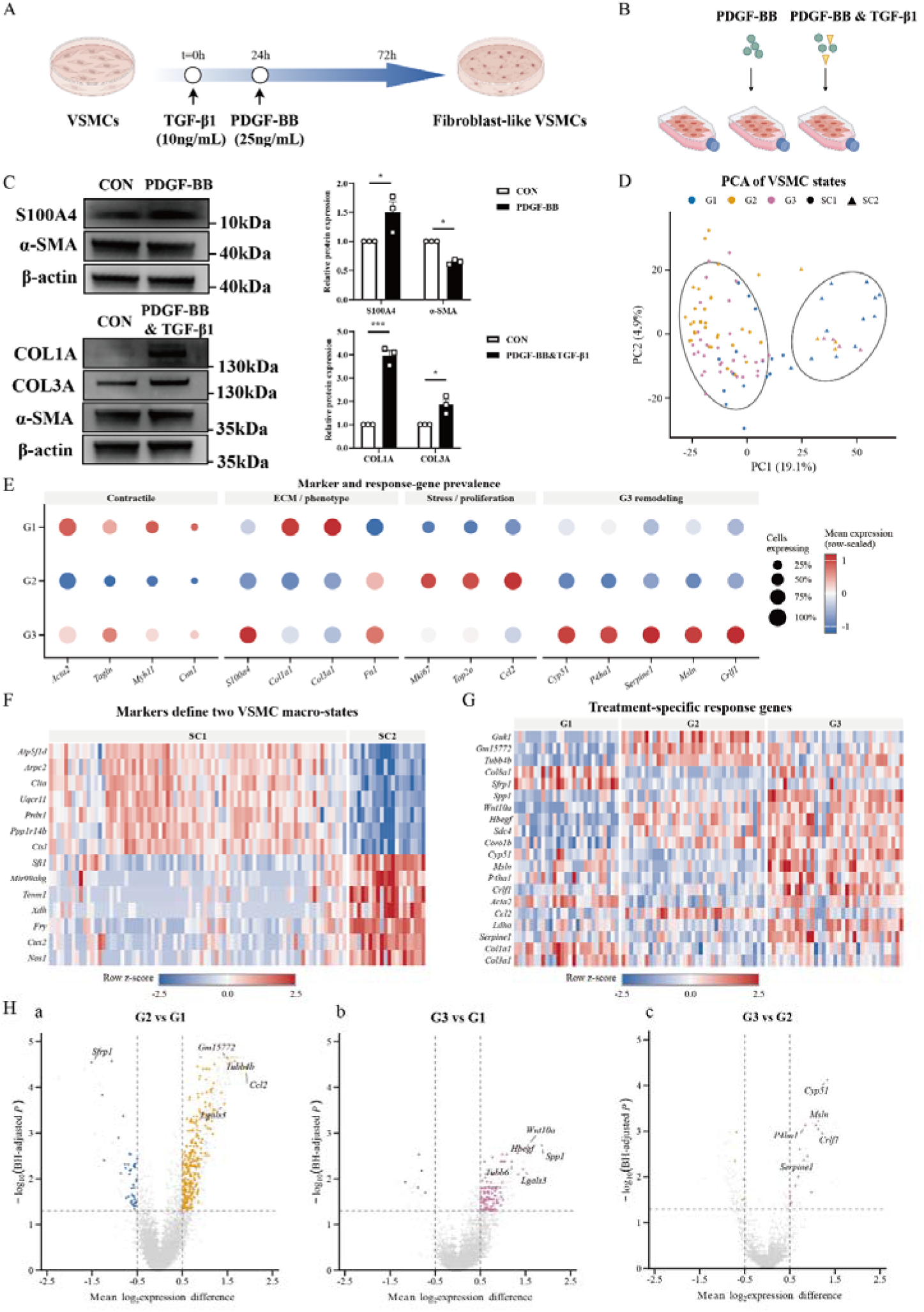
Combined stimulation partly restores contractile features and establishes a remodelling state in VSMCs. **A**, Experimental timeline. **B**, Treatment-group design. **C**, Representative western blots and quantification of S100A4, α-SMA, COL1A and COL3A; β-actin was the loading control. **D**, PCA of 99 primary-QC cells (G1, 27; G2, 37; G3, 35). Colours denote treatment, and neutral contours denote the group-blind SC1 and SC2 macro-states. **E**, Expression of contractile, ECM or phenotypic, stress or proliferative, and G3-remodelling genes. Dot size denotes the expressing-cell fraction, and colour denotes gene-scaled mean log1p(CP10k) expression. **F**, Single-cell heat map of SC1 and SC2 markers. **G**, Single-cell heat map of treatment-response genes. **H**, Exploratory cell-level differential-expression plots for G2 versus G1, G3 versus G1 and G3 versus G2. Cell-level statistics prioritise candidates and do not substitute for biological replication.

Western blotting supported the expected treatment response (Figure 3C). Relative to G1, G2 showed increased S100A4 (P=0.0245) and reduced α-SMA (P=0.0422), indicating a shift in smooth muscle cells from a contractile to a synthetic phenotype. In G3, α-SMA showed partial recovery relative to G2, while COL1A and COL3A increased (P=0.0002 and P=0.0167, respectively), indicating a transition of smooth muscle cells from a contractile to a fibroblastic phenotype. The incomplete agreement between collagen protein and endpoint mean mRNA is compatible with differences in transcriptional timing, translation, and extracellular deposition.

Principal-component analysis of 3,000 highly variable genes showed a treatment-associated shift without complete separation of the three groups (Figure 3D). Group-blind clustering identified two broad expression states rather than three treatment-defined clusters: SC1 contained 79 cells and SC2 contained 20 cells. SC1 represented 55.6% of G1 cells, 89.2% of G2 cells, and 88.6% of G3 cells. The similar SC1/SC2 composition of G2 and G3 indicated that their molecular differences arose primarily within a shared broad state rather than from different proportions of discrete cell types. PCA, UMAP, and contractile, synthetic/extracellular-matrix (ECM), and proliferative module scores supported a continuous state structure (Supplementary Figure S9). Marker expression and within-state module scores further showed that treatment effects reflected both state composition and programme activity within SC1 or SC2 (Figure 3F; Supplementary Figure S10). Because SC2 contained only four G2 and four G3 cells, comparisons within this state were treated as descriptive.

Representative genes separated the PDGF-BB response from the effect of TGF-β1 pretreatment (Figure 3E, G). G2 cells showed reduced Acta2, Tagln, and Myh11, together with increased Mki67, Top2a and Ccl2. In G3, contractile markers shifted partly towards G1, and proliferative or stress-associated signals were attenuated. G3 also showed broad detection of Cyp51, P4ha1, Serpine1, Msln and Crlf1, forming a linked metabolic and tissue-remodelling programme. Exploratory cell-level differential-expression analysis identified 341 candidates in G2 versus G1, 117 in G3 versus G1 and 17 in G3 versus G2 (Figure 3H). The G2 response included higher Ccl2, Tubb4b and Lgals3 and lower Sfrp1. G3 versus G1 highlighted Spp1, Wnt10a and Hbegf, while G3 versus G2 prioritised Cyp51, Crlf1, Msln, P4ha1 and Serpine1.

A continuous expression-state axis initially correlated strongly with ribosomal-transcript fraction (Spearman ρ=0.918; Supplementary Figure S11). After linear adjustment for UMI depth, detected-gene number, mitochondrial and ribosomal fractions, and technical library, correlations with these quality variables approached zero. The adjusted axis retained dynamic expression of Pcolce, P4hb, and Cox4i1 (Supplementary Figure S12). Its median value was 0.454 in G1, 0.316 in G2 and 0.415 in G3, with G3 shifted towards G1 relative to G2 (exploratory adjusted P=0.0159). This result supports partial correction of the PDGF-associated state but does not establish a temporal trajectory or lineage transition.

### TGF-β1 pretreatment establishes a metabolic–matrix expression programme

Pathway and programme analyses resolved the additional G3 response (Figure 4). KEGG analysis associated G2 versus G1 with ribosome, oxidative phosphorylation, proteasome, DNA replication and spliceosome pathways. G3 versus G1 additionally highlighted carbon metabolism, glycolysis, amino-acid biosynthesis, mitophagy and HIF-1 signalling, while G3 versus G2 prioritised steroid biosynthesis, glycolysis, HIF-1, AMPK, PI3K–Akt and focal-adhesion pathways (Figure 4A). A quality-adjusted response map separated the initial PDGF-BB effect from the added effect of TGF-β1 pretreatment (Figure 4B). Thirty genes met the prespecified criteria for G3-specific upregulation, three showed recovery after PDGF-associated reduction, one showed reversal after PDGF-associated induction and 11 remained reduced. Acta2, Igfbp7 and Ifitm3 showed partial recovery; Ccl2 was induced by PDGF-BB and reversed in G3; and Spp1, Ldha, Pkm, Pgk1, Pgam1, Cyp51, Hmgcr, Lgals3 and Serpine1 were concentrated in the G3-specific region.

**Figure 4:**
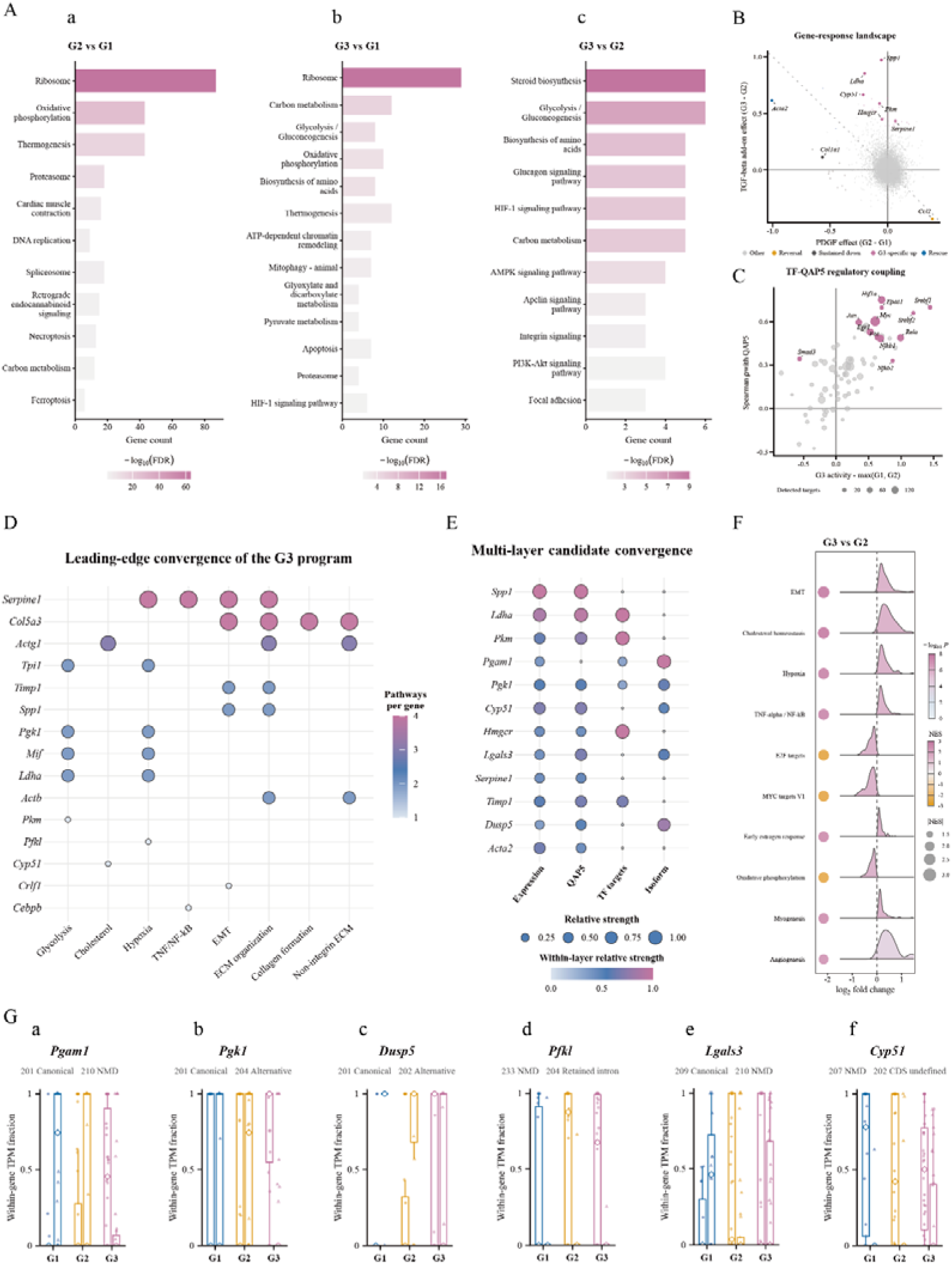
G3-specific metabolic–matrix programme links regulatory activity to transcript usage. **A**, KEGG enrichment of upregulated genes in the three pairwise treatment contrasts. **B**, Quality-adjusted gene-response map; the horizontal axis represents the G2−G1 effect, and the vertical axis represents the added G3−G2 effect. **C**, G3-associated inferred transcription-factor activity and correlation with QAP5. **D**, Leading-edge genes shared across enriched G3-versus-G2 pathways. **E**, Convergence of the G3−G2 expression effect, QAP5 loading, candidate transcription-factor targeting, and transcript-usage change. **F**, Hallmark GSEA ridge-bubble plot for G3 versus G2; bubble size indicates −log10(FDR). **G**, Single-cell within-gene TPM usage for two representative transcripts from *Pgam1*, *Pgk1*, *Dusp5*, *Pfkl*, *Lgals3* and *Cyp51*.

Non-negative matrix factorisation identified co-expression programmes without relying on predefined markers (Supplementary Figure S13). After adjustment for quality covariates, five quality-adjusted programmes (QAP1–QAP5) showed reduced correlations with sequencing complexity. QAP3, which included Ccl2 and stress-associated genes, was lower in G3 than G2 (adjusted P=4.32×10□□). QAP5 contained Spp1, Ldha, Pkm, Cyp51, Msmo1, Lgals3 and Crlf1 and was higher in G3 than both G1 and G2 (adjusted P=2.48×10□□and 1.55×10□□, respectively). Hallmark and Reactome gene-set enrichment analysis placed G2 versus G1 along MYC, E2F, oxidative-phosphorylation, DNA-repair and G2/M-checkpoint programmes (Supplementary Figure S14). By contrast, G3 versus G2 was positively enriched for epithelial–mesenchymal transition, cholesterol homeostasis, hypoxia, TNF-α/NF-κB signalling, myogenesis, angiogenesis and glycolysis, and negatively enriched for oxidative phosphorylation, MYC, E2F and cell-cycle programmes (Figure 4F; Supplementary Figure S15A).

Inferred transcription-factor activity linked SREBF1/2, HIF1A/EPAS1, NF-κB and AP-1 family regulators to QAP5 (Figure 4C; Supplementary Figure S15B). Leading-edge analysis identified Serpine1, Col5a3, Actg1, Ldha, Pgk1, Spp1 and Timp1 as shared contributors to several G3-versus-G2 pathways (Figure 4D). Integrating expression effects, QAP5 loading, candidate transcription-factor targeting, and transcript-usage changes prioritised Spp1, Ldha, Pkm, Hmgcr, Cyp51, Pgk1, and Pgam1 (Figure 4E). Changes in the fraction of expressing cells provided a complementary view of these responses (Supplementary Figure S15C). Together, G3 partly restored contractile features and suppressed the Ccl2/QAP3 stress response, while establishing a QAP5-dominated metabolic–ECM state. Persistent Spp1, Serpine1 and ECM signals also indicate a potential remodelling or fibrotic component; the G3 state therefore cannot be equated directly with clinical protection.

### Treatment-associated remodelling extends to transcript usage

Transcript-level analysis extended this response beyond gene abundance (Figure 4G; Supplementary Figure S16). Among 27,052 annotated transcripts, effect-size thresholds identified 996 G3-specific increases, 379 G3-specific decreases, 290 candidates that recovered after a PDGF-associated decrease and 209 candidates that reversed after a PDGF-associated increase. Gene-level transcript-usage effects derived independently from NumReads and TPM were strongly correlated across all three comparisons (Spearman ρ=0.933–0.938), with 89.5–90.3% directional agreement. Among 6,900 genes with at least two active transcripts, stringent abundance, detection, metric-concordance and technical-library filters yielded 414 descriptive candidates from 373 genes; 178 genes showed a change in the dominant annotated transcript. The candidates included 232 protein-coding and 142 nonsense-mediated-decay transcripts.

Pgam1, Pgk1, Dusp5, Pfkl, Lgals3 and Cyp51 illustrated coordinated transcript redistribution (Figure 4G). G3 increased Pgam1-201 while reducing the nonsense-mediated-decay transcript Pgam1-210; Pgk1-201 increased relative to Pgk1-204; and Dusp5-201 and Dusp5-202 showed near-mirror changes. Pfkl, Lgals3, and Cyp51 also redistributed usage among protein-coding, nonsense-mediated-decay, or retained-intron transcripts. These genes connect glycolysis, MAPK feedback, inflammatory or matrix remodelling, and cholesterol metabolism, providing orthogonal support for the gene-and pathway-level findings. The analysis estimates relative usage of reference-annotated transcripts and does not by itself validate splice junctions. Transcript-specific long-read sequencing will be required to confirm the candidate isoforms.

## Discussion

CHART-seq addresses a practical bottleneck in full-length single-cell RNA sequencing by moving sample identification to the heteroduplex-tagmentation step. Orthogonal Tn5 barcode combinations allow pooling wells before purification, gap filling, and PCR, replacing many parallel libraries with one shared library. In the tested workflow, this architecture shortened preparation to less than 3 h and reduced reagent cost to below US<$>1 per cell, while retaining UMI-based molecule counting and broad transcript coverage. The principal advantage is therefore not a single biochemical improvement, but a change in when samples become traceable and can safely be combined(16,17).

The matched-depth benchmark indicates that this simplification did not require a major loss of sensitivity. CHART-seq detected more genes and annotated isoforms than the comparator datasets and showed high cell-to-cell concordance. Its relatively high intronic fraction, together with a low intergenic fraction, is compatible with capture of incompletely spliced RNA rather than nonspecific genomic background(8,33). That feature could improve sensitivity to recent transcriptional responses. However, the benchmark used data generated by different studies and cannot separate chemistry from cell handling, laboratory practice, sequencing configuration, or computational preprocessing. A definitive comparison will require the same cell suspension to be split across methods, processed in parallel, sequenced on the same run, and analysed with matched molecule definitions(3–6,35,36).

The VSMC experiment illustrates the type of biological question enabled by the method. PDGF-BB shifted cells away from a contractile profile and towards proliferative and inflammatory expression. TGF-β1 pretreatment partly restored contractile features and reduced the Ccl2-associated QAP3 programme, consistent with attenuation of one component of the PDGF-BB response. The combined treatment did not simply return cells to the untreated state(37–40). It induced QAP5, which linked glycolysis, cholesterol metabolism, hypoxia-associated regulation and ECM remodelling, and this signal was supported by pathway enrichment, inferred regulatory activity and transcript-usage changes. The convergence of SREBF, HIF, NF-κB and AP-1-associated signals provides a testable model in which metabolic adaptation and matrix remodelling are coordinated rather than independent responses(41–45).

This interpretation also places a boundary on the clinically favourable framing of G3. Recovery of α-SMA and reduction of proliferative or Ccl2-associated signals may indicate a more stable VSMC phenotype than G2. Conversely, increased collagen proteins, Spp1, Serpine1 and other ECM-associated genes could support repair in one context but fibrosis or maladaptive remodelling in another(37,38,46,47). The present experiment measured one endpoint in cultured cells and cannot determine vessel-level function, plaque stability, or long-term clinical benefit. Time-resolved experiments, matrix-deposition assays, migration and contraction measurements, and in vivo vascular-injury models are needed to distinguish adaptive repair from persistent fibrosis(47–50).

The following technical limitations should be considered for further applications of CHART-seq. The 6-nt UMI provides 4,096 possible sequences, which may approach saturation for high-abundance molecules, particularly under conditions of high RNA input or deep sequencing. Therefore, barcode conflicts, index skips, and cross-well contamination should be quantified using species mixtures and blank well controls(51,52). A slight 3′-end bias indicates that maintaining RNA integrity before labeling remains important; if the DNase I treatment step affects sequencing coverage, it should be optimized or replaced. SHERRY showed that introducing the D188E substitution into the hyperactive pTXB1 Tn5 background markedly impaired its dsDNA fragmentation activity(15). This mutation may therefore reduce the tagmentation of contaminating genomic DNA in CHART-seq and could potentially permit omission of the DNase I treatment step, provided that sufficient activity towards RNA–cDNA heteroduplexes is retained. This possibility requires experimental validation. Furthermore, short-read sequencing cannot directly validate complete transcripts; therefore, transcript-level results must be confirmed via adapter-specific long-read sequencing(31,32,52,53). Routine monitoring of synthetic support DNA is also necessary to ensure that it does not introduce alignable contaminants or affect the selection of fragment sizes(53–55).

In summary, CHART-seq combines early combinatorial indexing, direct RNA–cDNA heteroduplex tagmentation and UMI-based counting in a rapid pooled workflow. The method offers high gene and transcript recovery at modest sequencing depth and can resolve coordinated gene-programme and transcript-usage responses. With direct same-sample benchmarking, expanded collision controls and independent biological validation, CHART-seq could provide a practical platform for scalable full-length single-cell transcriptomics.

## Materials and Methods

### Animals

All animal experiments conducted in this study strictly adhered to the principles outlined in the “Guide for the Care and Use of Laboratory Animals” as published by the National Academy of Sciences and the National Institutes of Health. The animal protocols and procedures were granted approval by the Animal Care and Use Committee of Nanjing First Hospital, Nanjing Medical University (DWSY-23151455). Furthermore, all animals were housed in a temperature-controlled environment with a 12-hour light/dark cycle and provided with free access to fresh water and food. In this study, the C57BL/6J mice were acquired from Cyagen Biosciences.

### Cell cultures

The HepG2 cell line was purchased from ATCC and incubated at 37 °C with 5% CO2 in Dulbecco’s modified Eagle medium (DMEM) (Gibco, 11965092), which was supplemented with 10% fetal bovine serum (FBS) (Gibco, 1600044) and 1% penicillin-streptomycin (Gibco, 15140122). Cells were dissociated with 0.05% Trypsin-EDTA (Gibco, 25300062) at 37 °C for 4 min and washed with D-PBS (STEMCELL, 37350).

HEK 293T cell line was purchased from the Cell Bank of the Chinese Academy of Sciences. and incubated at 37 °C with 5% CO2 in culture medium specifically formulated for HEK-293T cells (iCell, iCell-h237-001b).

Primary mouse vascular smooth muscle cells (VSMCs) were isolated from the thoracic aortas of 10-week-old male mice (C57BL/6J) by collagenase digestion. Briefly, the thoracic aorta was isolated, cleaned of endothelial cells and adventitial tissue, and incubated with 400U/ml collagenase II and 2.5U/ml elastase in DMEM at 37 for 1.5 h. The cell suspension was washed with complete medium, filtered through a 70-μm strainer, and plated onto 1 well of a 6-well culture plate. The cells were allowed to adhere for 3 days before replacing the medium. Isolated VSMCs were maintained in low-glucose Dulbecco’s modified Eagle medium containing 10% fetal bovine serum, and cells at passage 3 were used for further experiments.

### RNA extraction

After removing the cell culture medium, add 5 mL of Trypsin-EDTA (0.05%) (STEMCELL, 07910) to digest the cells for 10 min. After digestion, transfer the cell suspension to a sterile centrifuge tube and centrifuge at 600 × g for 5 min at 4 °C to remove trypsin. The cells were then washed twice with 5 mL of D-PBS, with centrifugation at 600 × g for 5 min at 4 °C after each wash. After washing, 1 mL of TRIzol Reagent (Invitrogen, 15596018CN) was added, and the sample was pipetted up and down and incubated at room temperature for 5 min. Subsequently, 100 µL of chloroform substitute (Servicebio, G3014-01) was added. The tube was capped tightly and vigorously shaken for 30 s until the mixture became turbid, followed by incubation at room temperature for 5 min. The sample was then centrifuged at 10,000 × g for 10 min at 4 °C. The upper colorless aqueous phase was carefully transferred to a new centrifuge tube, and an equal volume of isopropanol (Aladdin, I112021) was added. The sample was mixed by inversion and incubated at room temperature for 20 min. After centrifugation at 10,000 × g for 10 min at 4 °C, the supernatant was discarded. The pellet was washed twice with 1 mL of 75% ethanol, with centrifugation at 10,000 × g for 3 min at 4 °C after each wash. The supernatant was discarded, and the pellet was air-dried at room temperature for 5 min. An appropriate volume of TE Buffer (Sangon Biotech, B548408-0500) was then added to fully dissolve the RNA. RNA concentration was measured using an RNA rapid quantification kit suitable for Qubit (BBI, N608303) on a Qubit 4.0 fluorometer. The RNA concentration was ensured to be above 10 ng/µL, and the samples were stored at −80 °C.

### Combinatorial Barcode Tn5 Transposase Preparation

The bare Tn5 transposase used in this study was purchased from Accurate Biology (AG, AG12513). All primers for this study were purchased from Sangon Biotech (Shanghai), and the information is provided in Supplementary Table S1, in which the ME sequence is the chimeric end of the Tn5 transposon, and the sequence design of ME-A_m_ and ME-B_n_ (5′ to 3′) is described below: 1) The sequence that forms a reverse-complementary binding with the ME sequence; 2) an eight-base barcode A_m_/B_n_ index was used to label fragmented dsDNA fragments. We have listed twenty different barcodes in this study, which can be used to label up to 400 different samples by orthogonal combination barcoding, and the barcode can be further flexibly increased according to the experimental requirements of higher throughput without other conditions changing; 3) UMI was available only for ME-B_n_, which consists of a 6-nt concatenated base N for precise quantitative sequencing of expression levels of reads; 4) Read 1 and Read 2 were served as binding sites for sequencing primer recognition.

First, 4 µL ME-A_m_ (20 µm), 4 µL ME (20 µm), and 2 µL annealing buffer (Abclonal, RM20821) were added to a tube, vortexed, centrifuged, and collected at the bottom of the tube, then placed in a PCR thermocycler for hybridization to obtain the Mix A_m_ product. The following program was used: 95 °C for 5 min, decrease by 0.1°C per second to 25°C, 25 °C for 30 min, and 4 °C hold. The protocol for Mix B_n_ preparation is the same as that described above for Mix A_m_, with the minor difference of replacing the ME-A_m_ sequence with ME-B_n_. Next, 5.5 µL of Mix A_m_/Mix B_n_, 5 µL Tn5 transposase (0.5 µg µL^−1^), 8.75 µL assemble buffer (Abclonal, RM20187), and 5.75 µL nuclease-free water were mixed and incubated in a PCR thermocycler at 25 °C for 1.5 h to complete the assembly of Barcode Tn5 transposase (For long-term storage, the glycerol concentration must be adjusted to 50%). The products were named Barcode A_m_ Tn5 and Barcode B_n_ Tn5, respectively. Am and Bn are numbered according to the barcode chosen. Please note that the Tn5 transposon prepared using this protocol has a concentration of 100 ng/μL, which is suitable for CHART-seq with a 1 ng starting amount. If you need to perform CHART-seq with a lower starting amount, first prepare a Tn5 transposon solution at least 10 ng/μL, then serially dilute it to the desired concentration.

### CHART-seq library construction

For HEK293T cells, cells were digested with trypsin, washed twice with D-PBS, and filtered through a 70 μm sterile strainer. Subsequently, cells were stained using an Annexin V-APC/PI Apoptosis Detection Kit (Vazyme, A214), and double-negative cells were sorted on a BD FACSAria Fusion flow cytometer within 1 h after staining. For VSMCs, no staining was required; single cells were aspirated under a microscope using a laboratory-developed microneedle micromanipulator and transferred into PCR tubes(56).

For scRNA-seq library preparation of CHART-seq, single cells were sorted into 96-well plates containing 1.5 μL of lysis buffer composed of 1× Lysis Buffer (TAKARA, 635013), 0.3 U Recombinant RNase Inhibitor (TAKARA, 2313A), and 0.3 U AG DNase I (Thermo, 18068015). The plates were immediately centrifuged and incubated at 25 °C for 15 min for DNA digestion, followed by DNase I inactivation at 65 °C for 10 min. The plates were either stored at −80 °C or processed immediately. Subsequently, 0.5 μL of denaturation buffer containing 8 μM OligodTs (T30VN) and 8 mM dNTPs (BBI, A610057) was added, and the reaction was incubated at 72 °C for 3 min to facilitate RNA denaturation. Reverse transcription (RT) was then performed by adding 2 μL of RT mix (12 U SuperScript IV (Thermo, 18090050), 2× SSIV buffer, 10 mM DTT, 4 U RRI, and 2 M Betaine (Realtimes, RTE3103-02)), followed by incubation at 50 °C for 50 min and inactivation of the reverse transcriptase at 80 °C for 10 min. The resulting RNA/DNA hybrids, mixed with 1 μL of solution containing 100 pg DNA carrier, were tagmented with 0.5 μL Tn5 Am (400 pg/μL) and 0.5 μL Tn5 Bn (400 pg/μL) at 55 °C for 10 min (for 1 ng input samples, 200 ng each of Tn5 Am and Tn5 Bn were required), in a 4 μL reaction mix containing 2.5× TD buffer (25 mM Tris-HCl (Sangon Biotech, B548138-0500), 12.5 mM MgCl (BBI, B601192), 25% N,N-dimethylformamide (Macklin, N807509)), 22.5% PEG8000 (Beyotime, ST484), and 0.625 mM ATP (BBI, A600311). After the Tn5 reaction, 0.5 μL of Support DNA (2 ng/μL) was added to each sample, mixed by centrifugation, and all samples were collected into a single centrifuge tube. Purification was performed following the standard 5× protocol of DNA Clean & Concentrator-5 (ZYMO RESEARCH, D4014), with the following modifications: only 600 μL of Wash Buffer was used for a single wash, and elution was carried out with 20 μL of nuclease-free water (centrifugation at 16,000 g, 20 °C, 1 min). Subsequently, 0.25 μL of Bst 3.0 (NEB, M0374S) and 20 μL of Q5 high-fidelity 2× master mix (NEB, M0492L) were used to repair the gap left by Tn5 at 72 °C for 15 min, followed by termination of the reaction at 95 °C for 5 min. Finally, 2.5 μL of index I5 (10 μM), 2.5 μL of index I7 (10 μM), and 5 μL of Q5 were added. PCR was cycled as follows: initial denaturation at 98 °C for 30 s; 18 cycles of 20 s at 98 °C (only 9–12 cycles were required for 1 ng input), 20 s at 65 °C, and 2 min at 72 °C; and a final extension at 72 °C for 5 min. The indexed products were pooled and purified at 0.75× with VAHTS DNA Clean Beads (Vazyme, N411). The resulting libraries were sequenced on a NovaSeq 6000 or NovaSeq X Plus platform with PE150 configuration.

### Carrier DNA and Support DNA Preparation

Carrier DNA for tagmentation is produced by dissolving salmon sperm DNA (Aladdin, D404592) in 1× TE Buffer, quantifying it using the Equalbit 1× dsDNA HS Assay Kit (Vazyme, EQ121) on a Qubit 4.0 fluorometer, and diluting it to 100 pg/µL. Perform MDA amplification using EquiPhi29™ DNA polymerase (Thermo Scientific, A39390) as follows: mix 1 µL of the diluted salmon sperm DNA with 2 µL of 10× EquiPhi29™ DNA Polymerase Reaction Buffer, 0.2 µL of 0.1 M DTT, 2 µL of Exo-Resistant Random Primer (500 µM) (Thermo Scientific, SO181), 0.2 µL each of 100 µM dATP (BBI, A620046), dUTP (BBI, B600006), dGTP (BBI, A660046) and dCTP (BBI, A640046), and 13 µL of nuclease-free water. Gently vortex the mixture and briefly centrifuge to collect the contents at the bottom of the tube. Incubate the reaction mixture in a thermal cycler at 95 °C for 3 min, then immediately place the tube on ice. Add 1 µL of EquiPhi29™ DNA polymerase, mix well, and incubate in a PCR instrument at 42 °C for 2 h, followed by 65 °C for 10 min.

Support DNA for purification is produced by dissolving Support DNA1 and Support DNA2 in 1× TE Buffer to a concentration of 100 μM. Mix equal volumes of 100 μM Support DNA1 and Support DNA2, then place the mixture in a PCR instrument. The following program was used: 95 °C for 5 min, cooling at a rate of 0.1 °C per second to 25 °C, 25 °C for 30 min, and hold at 4 °C. Subsequently, store long-term at −80°C.

### Bioinformatics pipeline for CHART-seq

All downsampling was performed using seqtk. The raw sequencing data were stored in FASTQ format. First, FastQC was used for quality control of the raw data, after which umi-tools was used for cell barcode identification and extraction of valid reads. The specific parameters were as follows: --bc-pattern=“CCCCCCCC” --bc-pattern2=”CCCCCCCCNNNNNN” --set-cell-number=MIX_CELL_NUMBER --error-correct-threshold 0 --log2stderr. Subsequently, BBMap reformat was used to trim the 19-nt ME sequence. Cutadapt was then used to remove the adaptors AGATCGGAAGAG and CTGTCTCTTATACACATCT, as well as polyA/T/C/G stretches longer than 6 nt, retaining reads with a length of at least 20 bp and a quality score of at least 20. The trimmed FASTQ files were then aligned to the reference genome in paired-end mode using STAR. featureCounts was used to annotate and quantify the aligned reads, and umi-tools was finally used to integrate the gene expression matrix. For alignment-region statistics, Picard CollectRnaSeqMetrics was used, and for gene-body coverage calculation, geneBody_coverage.py from RSeQC was used.

### Data from other full-length RNA-seq methods

Flash-seq, Smart-seq3, and Smart-seq2 data were obtained from the NCBI Sequence Read Archive (accession no. PRJNA816486), and SHERRY2 data were obtained from the NCBI Sequence Read Archive (accession no. PRJNA879104). Detailed data accession numbers are provided in Supplementary Table S2.

For all datasets, 500K reads were downsampled with seqtk. Cutadapt was then used to remove the adaptors AGATCGGAAGAG and CTGTCTCTTATACACATCT, as well as polyA/T/C/G stretches longer than 6 nt, retaining reads with a length of at least 20 bp and a quality score of at least 20. The trimmed FASTQ files were then aligned to the reference genome in paired-end mode using STAR. featureCounts was used to annotate and quantify the aligned reads. The trimmed FASTQ files were aligned using Salmon; for single-end sequencing data, the additional parameters --fldMean 700 --fldSD 100 --fldMax 2000 were applied. SUPPA was used to quantify various alternative splicing (AS) events.

A gene was considered detected when its RPKM or FPKM exceeded 1, unless otherwise specified. For the five-method gene-overlap analysis, a gene was retained if its RPKM or FPKM was greater than 1 in at least 10% of the samples within a given method. Cell-to-cell reproducibility was summarized using pairwise Kendall’s τ after excluding the diagonal. For method comparisons, distributions were visualized using violin and box plots. CHART-seq was compared pairwise with each alternative method using a two-sided Mann–Whitney U test. Optimization experiments involving three or more conditions were analyzed using a Kruskal–Wallis test followed by Dunn’s post hoc test with Bonferroni correction, whereas two-condition experiments were analyzed using a two-sided Mann–Whitney U test.

### Statistics and Analysis

Statistical significance was defined as *P < 0.05, **P < 0.01 and ***P < 0.001.

### Western Blot

Whole-cell lysates were prepared using RIPA lysis buffer (Beyotime, P0013C) supplemented with 1× protease inhibitor (MCE, HY-K0010) and a phosphatase inhibitor cocktail (MCE, HY-K0021, HY-K0022). Equal amounts of total protein extracted from rats, murine tissues, or PASMCs were resolved by SDS-PAGE and subsequently transferred onto PVDF membranes (Bio-Rad). Following transfer, the membranes were blocked with 5% bovine serum albumin in Tris-buffer saline at 4°C overnight. Subsequently, membranes were probed with primary antibodies and then with horseradish peroxidase-conjugated secondary antibodies (Biosharp, BL001A, BL003A). Antibodies against S100A4 (16105-1-AP), α-SMA (14395-1-AP) and COL1A (14695-1-AP) were purchased from Proteintech Group, Inc. (Wuhan, China). Antibody against COL3A (A0817) was purchased from ABclonal, Inc. (Wuhan, China). Antibody against β-actin (AF7018) was purchased from Affinity Biosciences (Changzhou, China). The EasySee Western Blot Kit (TransGen, DW101-01) served as the HRP substrate for detection. Membranes were reprobed with another antibody after stripping using Re-Blot Plus Western Blot Mild Antibody Stripping Solution (Millipore, 2504). The EasySee Western Blot Kit (TransGen, DW101-01) served as the HRP substrate for detection. Membranes were reprobed with another antibody after stripping using Re-Blot Plus Western Blot Mild Antibody Stripping Solution (Millipore, 2504).

### VSMC expression-matrix quality control and normalisation

Gene-by-cell matrices contained UMI counts, with absent genes and zero-valued entries treated as not detected. For each cell, total UMIs, detected genes, mitochondrial-transcript fraction and ribosomal-transcript fraction were calculated. Relaxed quality control required at least 2,000 genes, at least 5,000 UMIs, and less than 25% mitochondrial counts. Stringent quality control required at least 3,000 genes, at least 10,000 UMIs, and less than 20% mitochondrial counts. The primary analysis additionally excluded complexity outliers, defined without group labels as values more than 3 median absolute deviations from the overall median on a log10(x+1) scale. Ninety-nine cells remained.

Counts were scaled to 10,000 UMIs per cell and transformed as ln[1+(gene UMI/total UMI)×10,000]. Among genes detected in at least three cells, mitochondrial and ribosomal genes were removed, and the 3,000 genes with the highest log1p(CP10k) variance were selected. Centred and scaled values were analysed by principal-component analysis using prcomp. Group labels were not used for state discovery. K-means solutions with k=2–6 were compared on PC1–PC10 using 200 random starts, and the selected k=2 model was refitted with 500 starts. Hierarchical clustering, alternative PC ranges, silhouette width and Gaussian-mixture modelling were used as sensitivity checks. UMAP used PC1–PC15 with cosine distance; n-neighbour values of 10, 15 and 25 and minimum-distance values of 0.10, 0.25 and 0.40 were compared, and the final setting maximised 10-neighbour trustworthiness rather than treatment separation.

### Module scores, differential expression and enrichment

VSMC identity, contractile, synthetic/ECM, proliferation, TGF-β-response and PDGF-response modules were defined from prespecified representative genes. Each gene was z-standardised across the 99 primary-analysis cells, and a module score was defined as the arithmetic mean of detected genes in the module. SC1/SC2 markers and group contrasts used two-sided Wilcoxon rank-sum tests with tie correction and rank-biserial effect sizes. Genes were tested if detected in at least 10% of either comparison group. Mean log2 expression difference was calculated as the difference in mean log1p(CP10k) expression divided by ln(2). P values were adjusted by the Benjamini–Hochberg (BH) method. Candidates required BH-adjusted P<0.05, an absolute mean log2 difference of at least 0.5, and an absolute difference of at least 0.15 in the fraction of expressing cells. Because cells were generated in a single experiment and technical libraries were not biological replicates, these cell-level P values were used only to rank exploratory candidates.

KEGG Mus musculus pathway definitions and gene mappings were obtained through KEGG REST. Upregulated candidates had BH-adjusted P<0.05 and mean log2 difference >0.5. The background comprised all tested genes that mapped to mouse KEGG identifiers. Pathways containing 10–500 background genes and at least three candidate genes were tested using a one-sided hypergeometric test with within-comparison BH correction. Disease-category pathways were omitted from the main display to reduce overinterpretation but retained in the source data. Preranked gene-set enrichment analysis used mouse Hallmark and Reactome collections from MSigDB v2026.1.Mm and fgseaMultilevel, with gene-set sizes of 15–500 and eps=1×10 . Genes were ranked principally by mean log2 difference, with small contributions from rank-biserial effect and detection-fraction difference to resolve ties. BH false-discovery rate (FDR)<0.05 defined significant enrichment.

### Continuous states, expression programmes and regulatory inference

A principal curve fitted to PC1–PC5 defined the unadjusted expression-state axis, which was scaled from 0 to 1 and oriented from SC2 to SC1. To reduce quality-related confounding, log1p(CP10k) expression was projected onto a design matrix containing log1p(total UMIs), log1p(detected genes), mitochondrial fraction, ribosomal fraction and technical library. A quality-adjusted state axis was fitted to PC1–PC5 of the residual matrix. Dynamic genes were ranked by Spearman correlation with the corresponding axis and adjusted by BH. LOESS curves used span=0.72 for visualisation only; the axes do not represent measured time, lineage or causality.

Non-negative matrix factorisation used the Brunet algorithm. Ranks 2–6 were assessed using five initialisations. Rank-2 and exploratory rank-5 models were then run 15 and 20 times, respectively. For quality-adjusted NMF, positive residual values were retained and a rank-5 model was run 15 times to obtain quality-adjusted programmes 1-5 (QAP1–QAP5). Programme activities were z-standardised across cells. Overall group differences were tested with Kruskal–Wallis tests, pairwise comparisons used Wilcoxon rank-sum tests with BH adjustment, and programme associations used Spearman correlation.

Quality-adjusted response categories were defined from mean residual-expression effects for G2−G1, G3−G2 and G3−G1. G3-specific increases required |G2−G1|<0.25, G3−G2≥0.35 and G3−G1≥0.35; decreases used the opposite signs. Recovery required G2−G1≤−0.35 and G3−G2≥0.35, whereas reversal required G2−G1≥0.35 and G3−G2≤−0.35. Welch’s t tests with BH adjustment were used only for auxiliary ranking. Mouse DoRothEA interactions were obtained through OmniPath. Only A–C confidence interactions with signed regulation were retained and weighted 1.0, 0.7 and 0.5, respectively. Transcription-factor activity was defined as the weighted signed mean of residual target-gene expression for factors with at least 10 detected targets, followed by cell-wise standardisation.

### Transcript abundance and within-gene usage

Transcript analyses used Salmon TPM and NumReads matrices together with transcript Length and EffectiveLength. Transcript identifiers were matched to a GTF-derived annotation containing gene identifier, gene symbol, transcript name, biotype, exon number, coding-sequence length and canonical status. Transcript-level structure was analysed from quality-and library-adjusted log1p(TPM) values. Within-gene usage was calculated separately from TPM and NumReads as the abundance of a transcript divided by the summed abundance of all annotated transcripts from the same gene in that cell or aggregate.

Active transcripts required at least 20 total NumReads and detection in at least 5% of primary-analysis cells. Candidate genes required at least two active transcripts, at least 200 gene-level NumReads, at least 50 transcript-level NumReads, at least 10% detection, a maximum NumReads-usage difference of at least 0.15, a concordant absolute TPM-usage difference of at least 0.10, and agreement in direction. High-confidence descriptive candidates further required at least 1,000 gene-level NumReads, at least 200 transcript-level NumReads, at least 20% detection, a maximum NumReads-usage difference of at least 0.20, an absolute TPM-usage difference of at least 0.15, and usage of at least 0.15 in one group. Technical-library consistency required directional agreement in at least two evaluable libraries. These filters identify annotation-supported usage candidates, not experimentally validated splice events.

### Software and versions

Java 18, python 3.12, R 4.6.0, FastQC v0.12.1, seqtk 1.3, umi-tools 1.1.6, cutadapt 5.0, STAR 2.7.11b, featureCounts v2.0.8, picard 3.3.0, rseqc 5.0.4, salmon 1.11.4, suppa 2.4 , matrix 1.7-5、matrixStats 1.5.0、ggplot2 4.0.3、ggrepel 0.9.8、patchwork 1.3.2、uwot 0.2.5、cluster 2.1.8.2、 mclust 6.1.3、fgsea 1.38.0、NMF 0.28、princurve 2.1.6、data.table 1.18.6.1、dplyr 1.2.1、scales 1.4.0、ragg 1.5.2 and svglite 2.2.2。

## Data Availability

Sequencing data related to HEK293T cells and HepG2 RNA have been deposited in the Sequence Read Archive (PRJNA1531701).

## Code Availability

Code and analysis scripts for this work are available on GitHub (https://github.com/DuDuDuDu-du/Chartseq_tools).

## Acknowledgements

This work was supported by the National Natural Science Foundation of China (81827901, 82361138570, 3260120387, 82602871), Jiangsu Provincial Medical Key Discipline (Laboratory) Cultivation Unit (JSDW202249).

## Author information

Wenyi Zhang and Aiqun Chen contributed equally to this work.

## Contributions

X.Z., X.G., and L.H. conceived the study. W.Z., A.C., and L.H. performed the experiments. W.Z. and J.S. performed the data analyses. Y.H., A.C., and D.Z. provided the samples. K.Y., H.W., Z.J., and Y.G. examined the manuscript. X.Z., Y. X., X.G., and L.H. supervised all aspects of this study. All authors read and approved the final manuscript.

---

**Figure S1:**
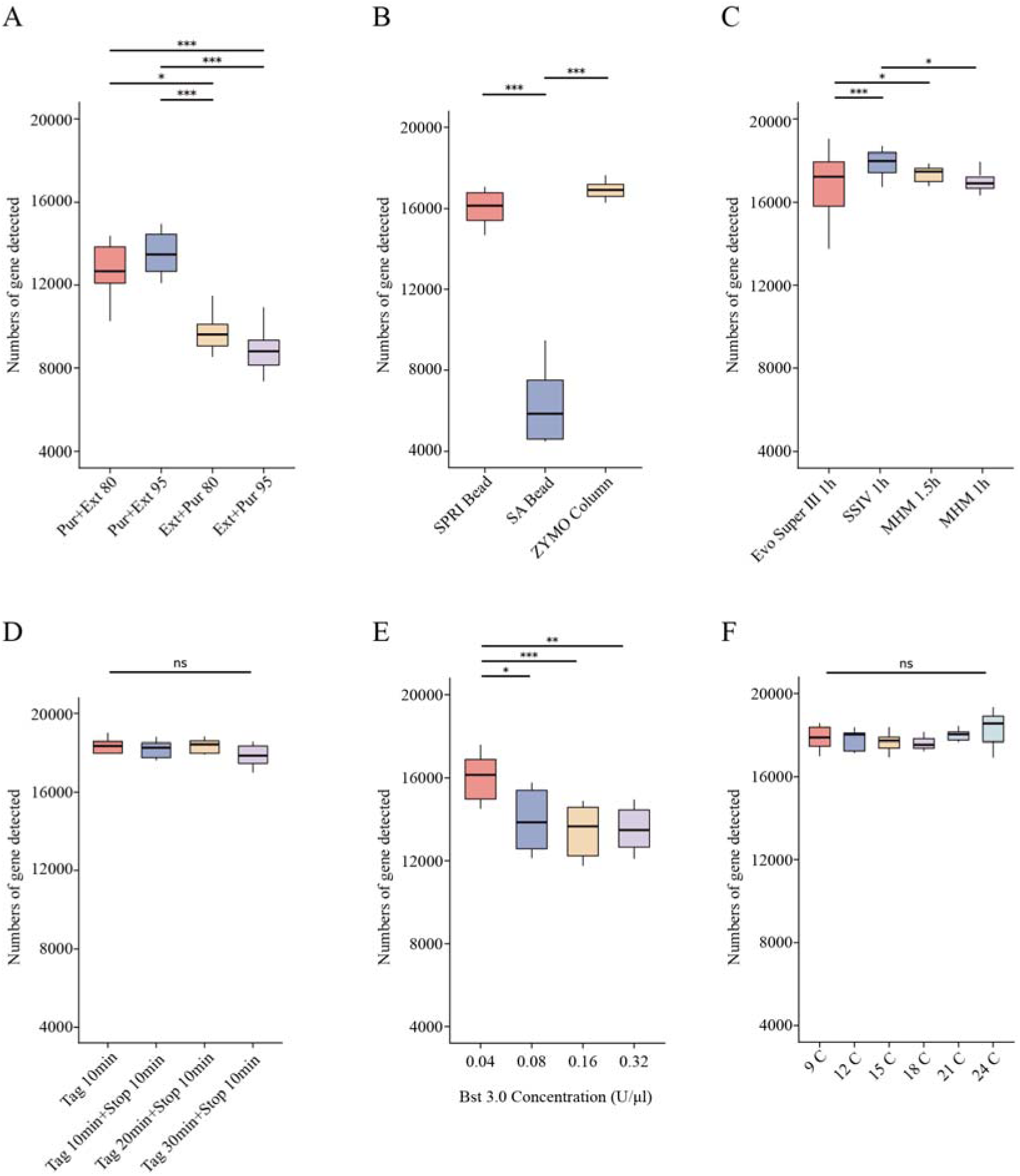
Establishment of the CHART-seq workflow at 1 ng RNA. Each condition comprised three pooled libraries with four RNA aliquots per library and was downsampled to 15 million reads per library. **A**, Order of purification and extension, and Bst 3.0 heat-inactivation temperature. **B**, Purification method. **C**, Reverse transcriptase and reaction duration. **D**, Tagmentation duration and heat stop. **E**, Bst 3.0 amount. **F**, PCR cycle number.

**Figure S2:**
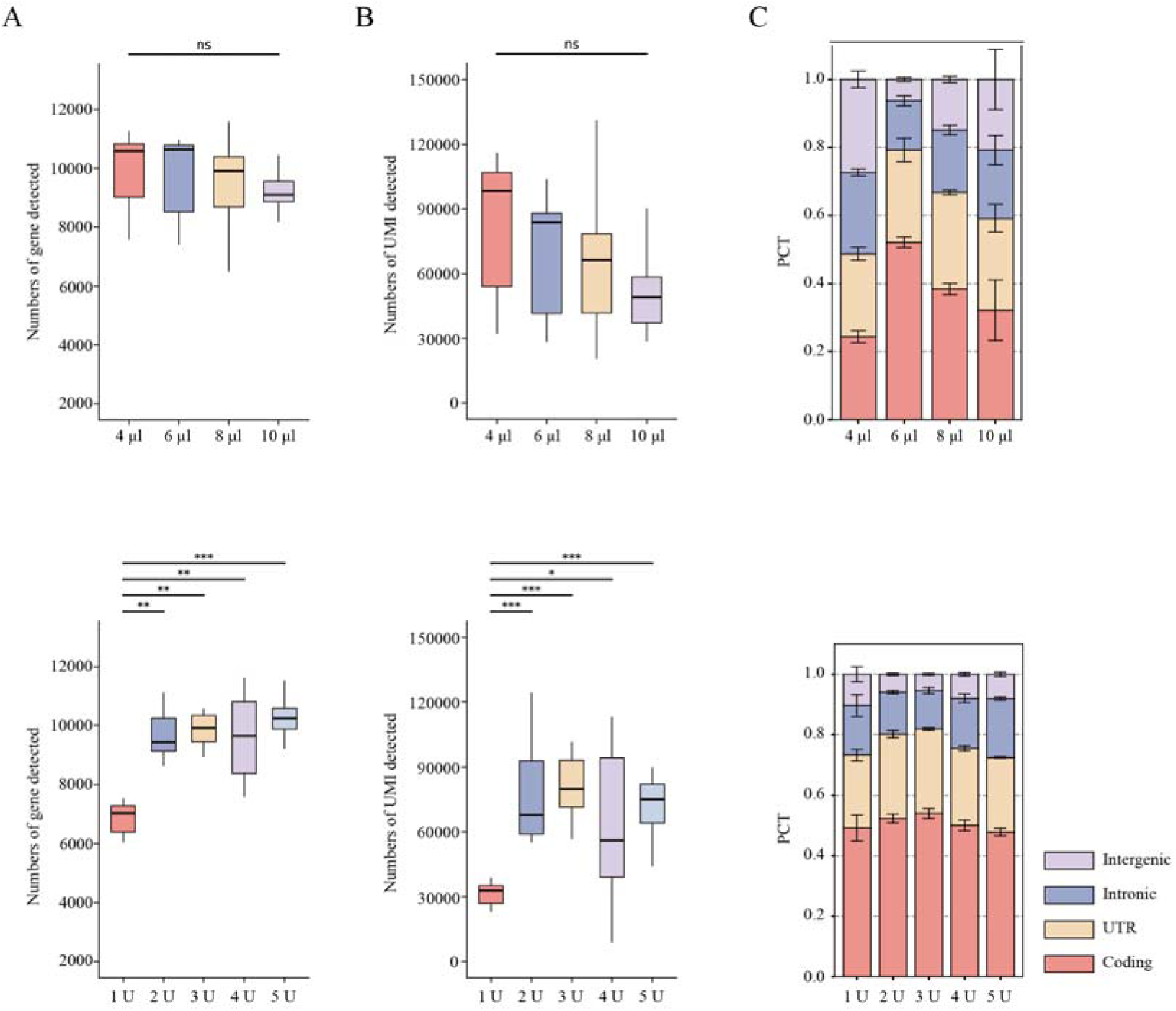
Optimisation of reverse transcription at 10 pg RNA. Each condition comprised three pooled libraries with four aliquots per library; libraries were downsampled to 5 million reads each. **A**, Detected genes. **B**, Detected UMIs. **C**, Genomic-region fractions across reverse-transcription volumes (top) and enzyme concentrations (bottom).

**Figure S3:**
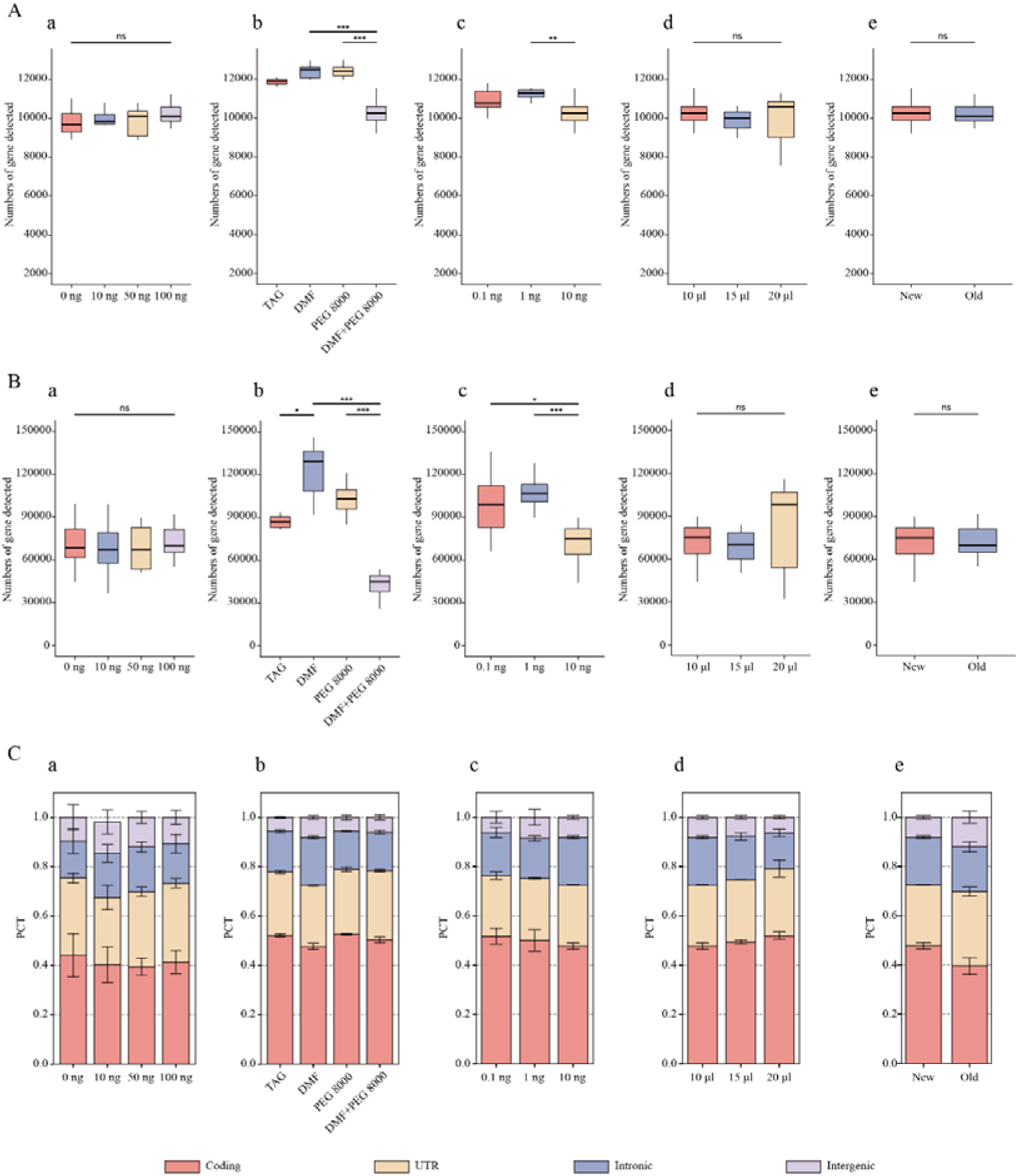
**Optimisation of heteroduplex tagmentation at 10 pg RNA**. **A**, Detected genes. **B**, Detected UMIs. **C**, Genomic-region fractions for carrier DNA (**a**), DMF and PEG 8000 (**b**), Tn5 amount (**c**), reaction volume (**d**) and buffer age (**e**). Replication and downsampling were as in Supplementary Figure S2.

**Figure S4:**
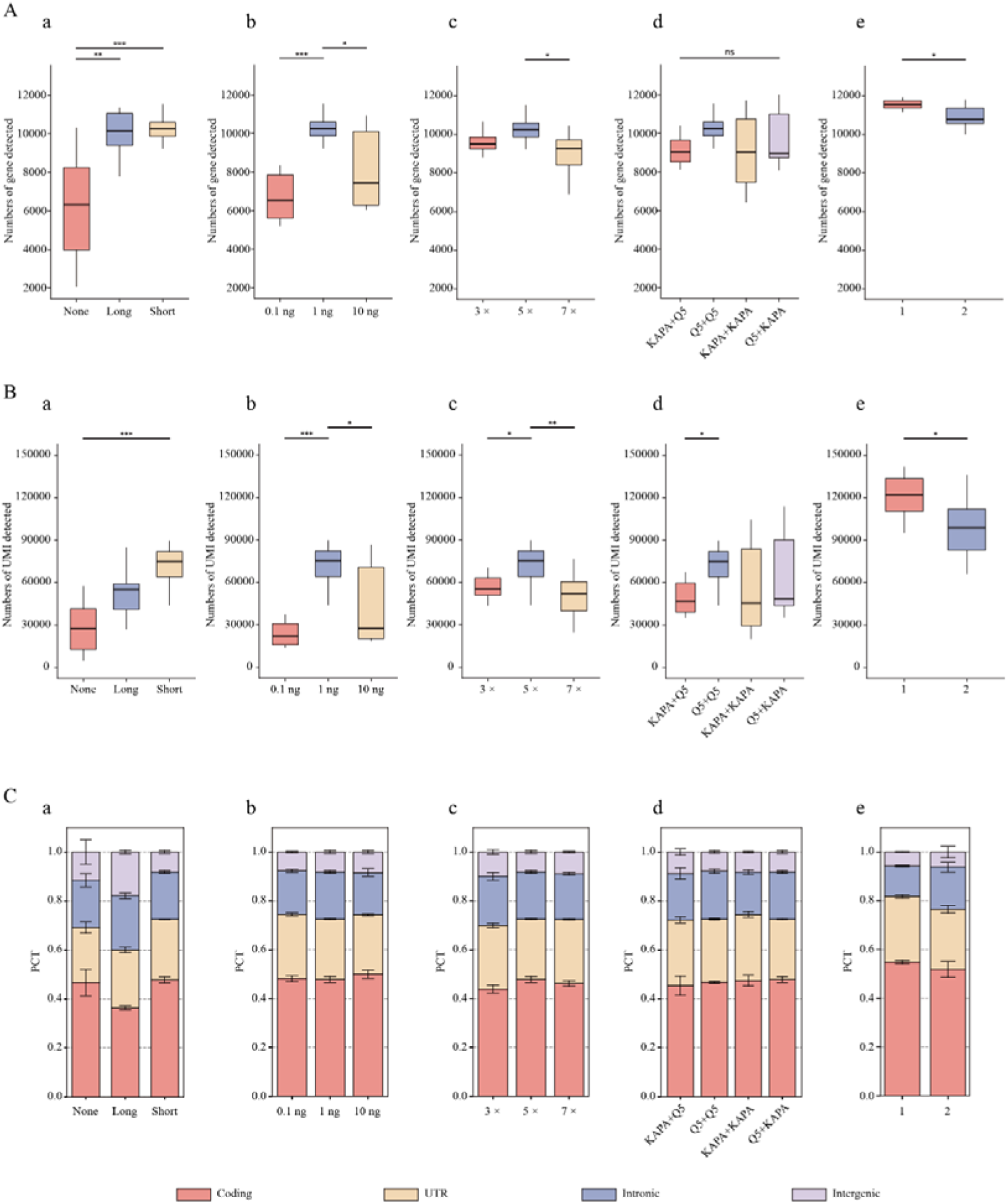
**Optimisation of pooled purification and library amplification**. **A**, Detected genes. **B**, Detected UMIs. **C**, Genomic-region fractions for support-DNA type (**a**), short support-DNA amount (**b**), binding-buffer amount (**c**), PCR enzyme (**d**) and number of purification steps (**e**). Replication and downsampling were as in Supplementary Figure S2.

**Figure S5:**
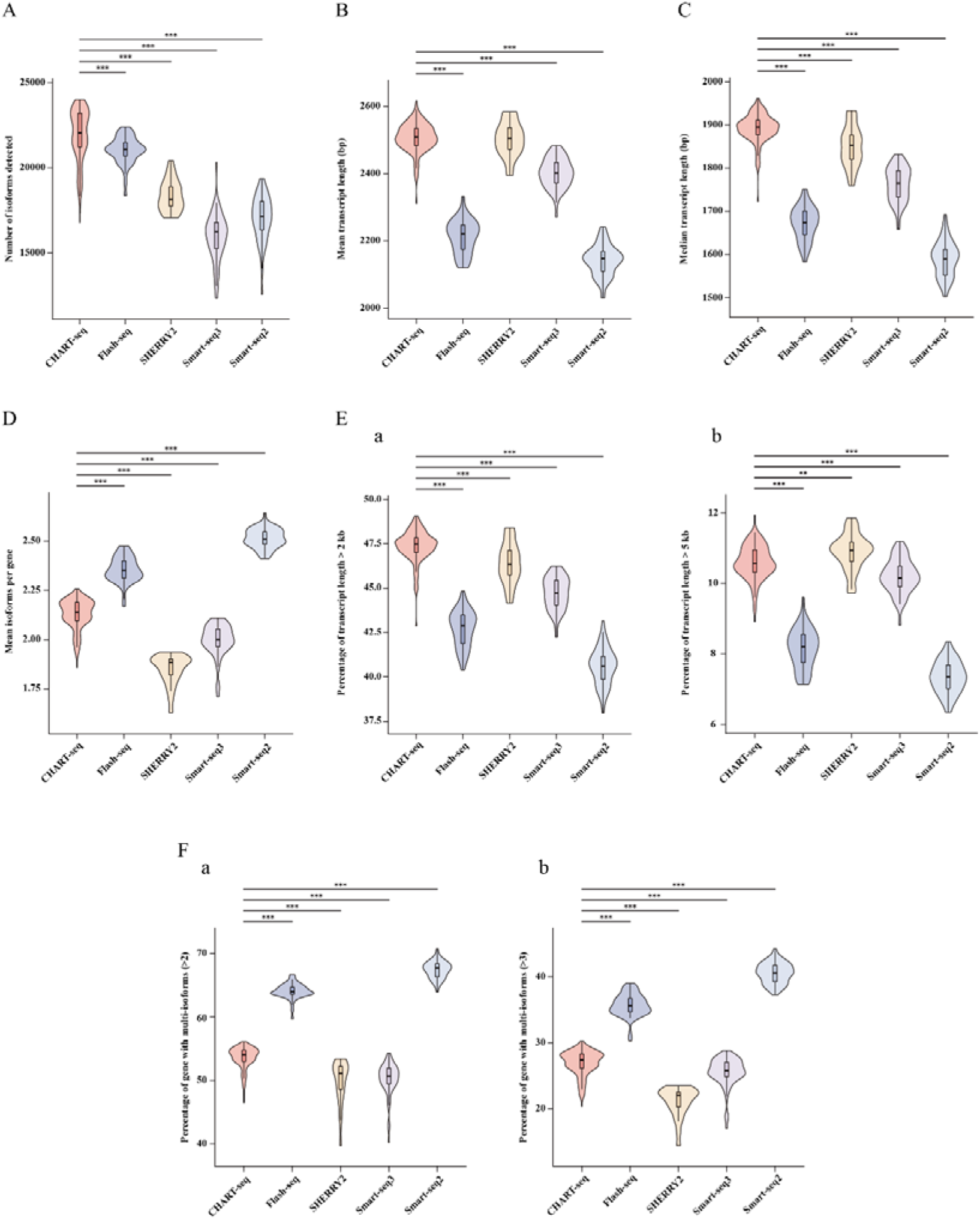
**Transcript-level benchmarking across full-length methods**. **A**, Detected isoforms. **B**, Mean detected-transcript length. **C**, Median detected-transcript length. **D**, Mean isoforms detected per gene. **E**, Fractions of detected transcripts longer than 2 kb (**a**) and 5 kb (**b**). **F**, Fractions of genes with at least two (**a**) or three (**b**) detected isoforms.

**Figure S6:**
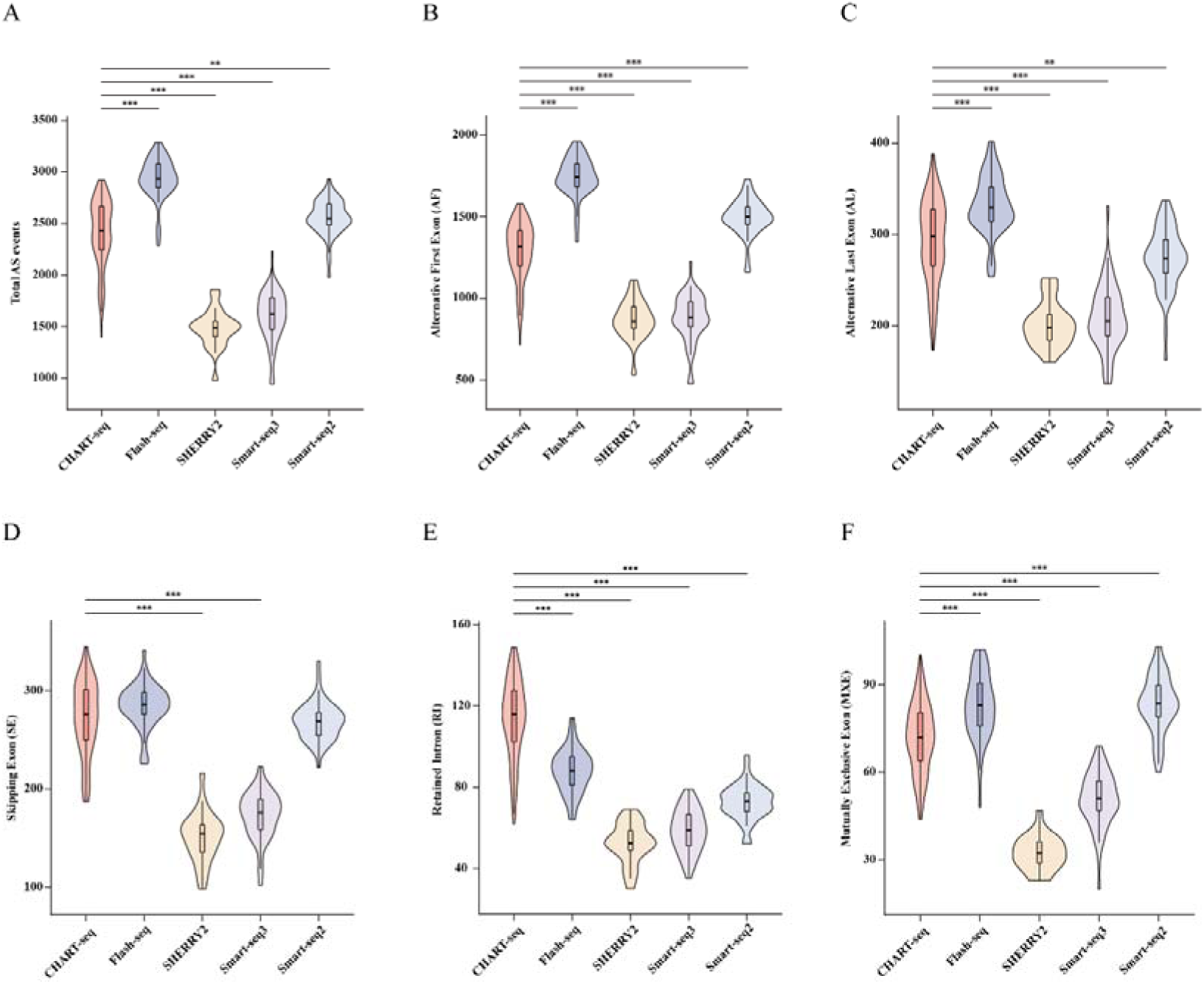
Alternative-splicing event detection across full-length methods. **A**, Total alternative-splicing events. **B–F**, Alternative first-exon, alternative last-exon, skipped-exon, retained-intron and mutually exclusive-exon events.

**Figure S7:**
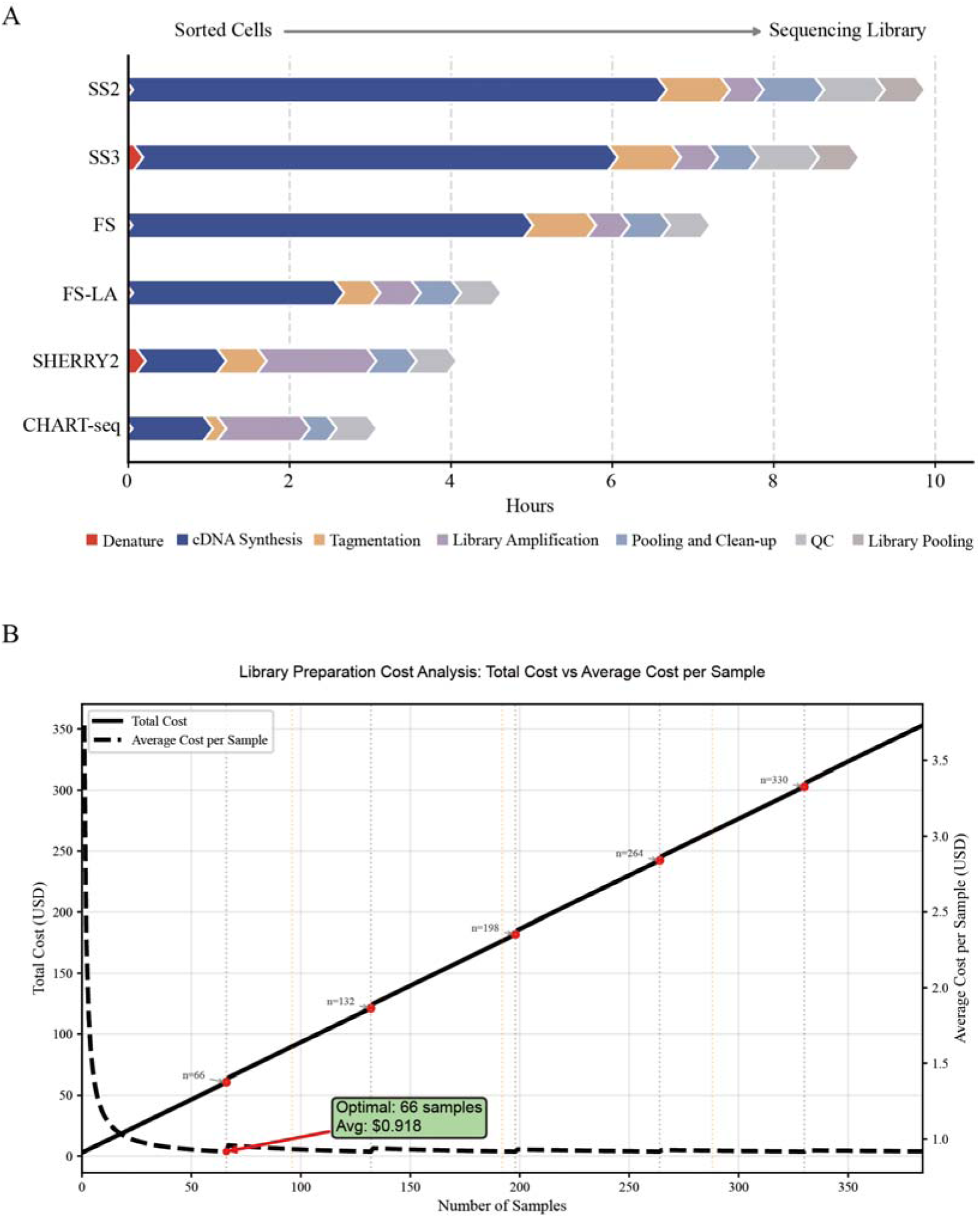
Library-preparation time and per-cell reagent cost. **A**, Preparation time from RNA denaturation to sequencing-ready libraries for five full-length methods. **B**, Estimated CHART-seq reagent cost per cell across pooling formats, calculated using an exchange rate of RMB 7 per US dollar.

**Supplementary Figure S8:**
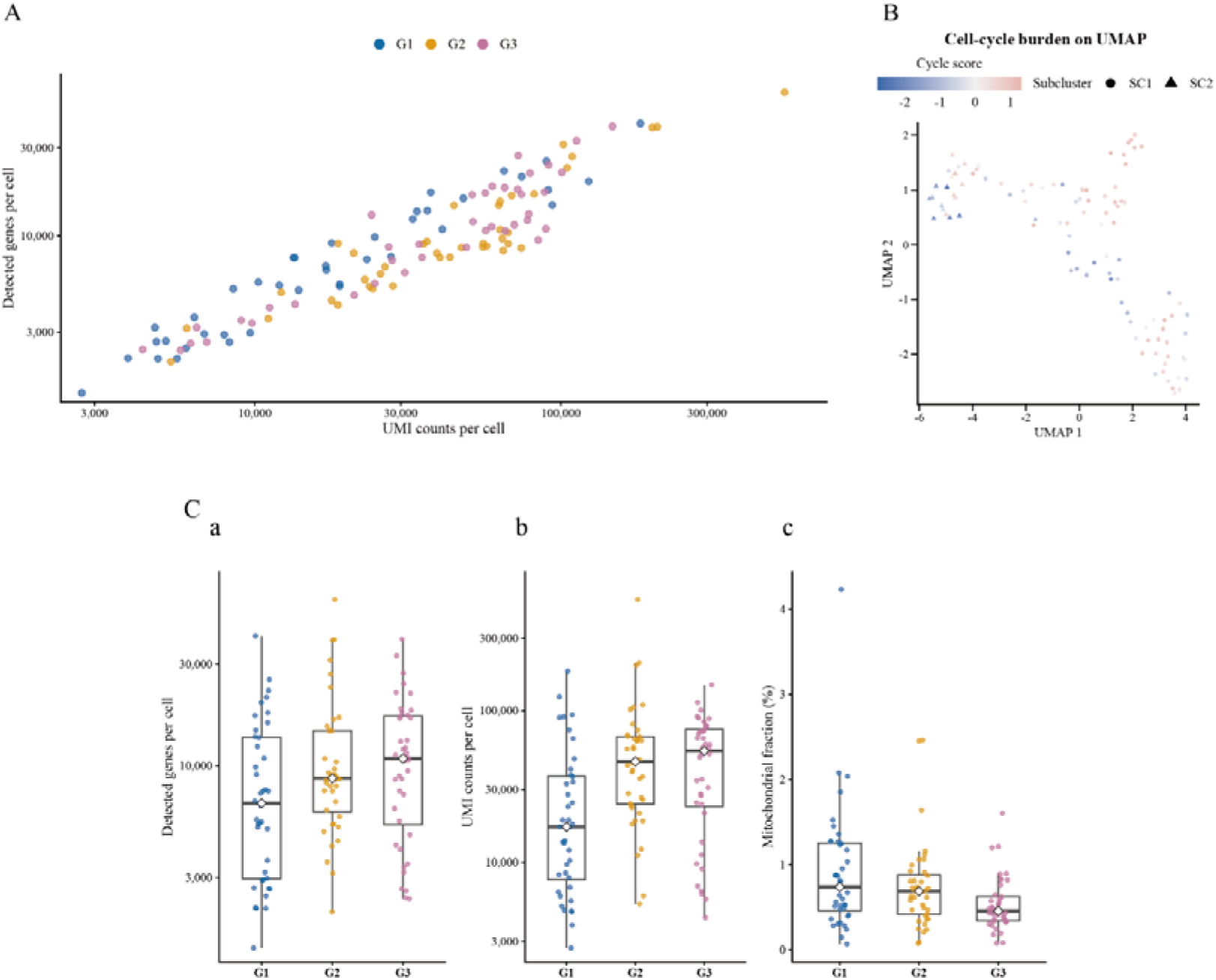
Quality assessment of the 120-cell VSMC design. **A**, Relationship between detected genes and total UMIs in the intended 120-cell design. Each group retained 14, 13 and 13 starting cells in B1, B2 and B3. **B**, Cell-cycle burden in the primary-QC UMAP, based on an E2F/G2M rank score. **C**, Distributions of detected genes (**a**), total UMIs (**b**) and mitochondrial-transcript fraction (**c**).

**Supplementary Figure S9:**
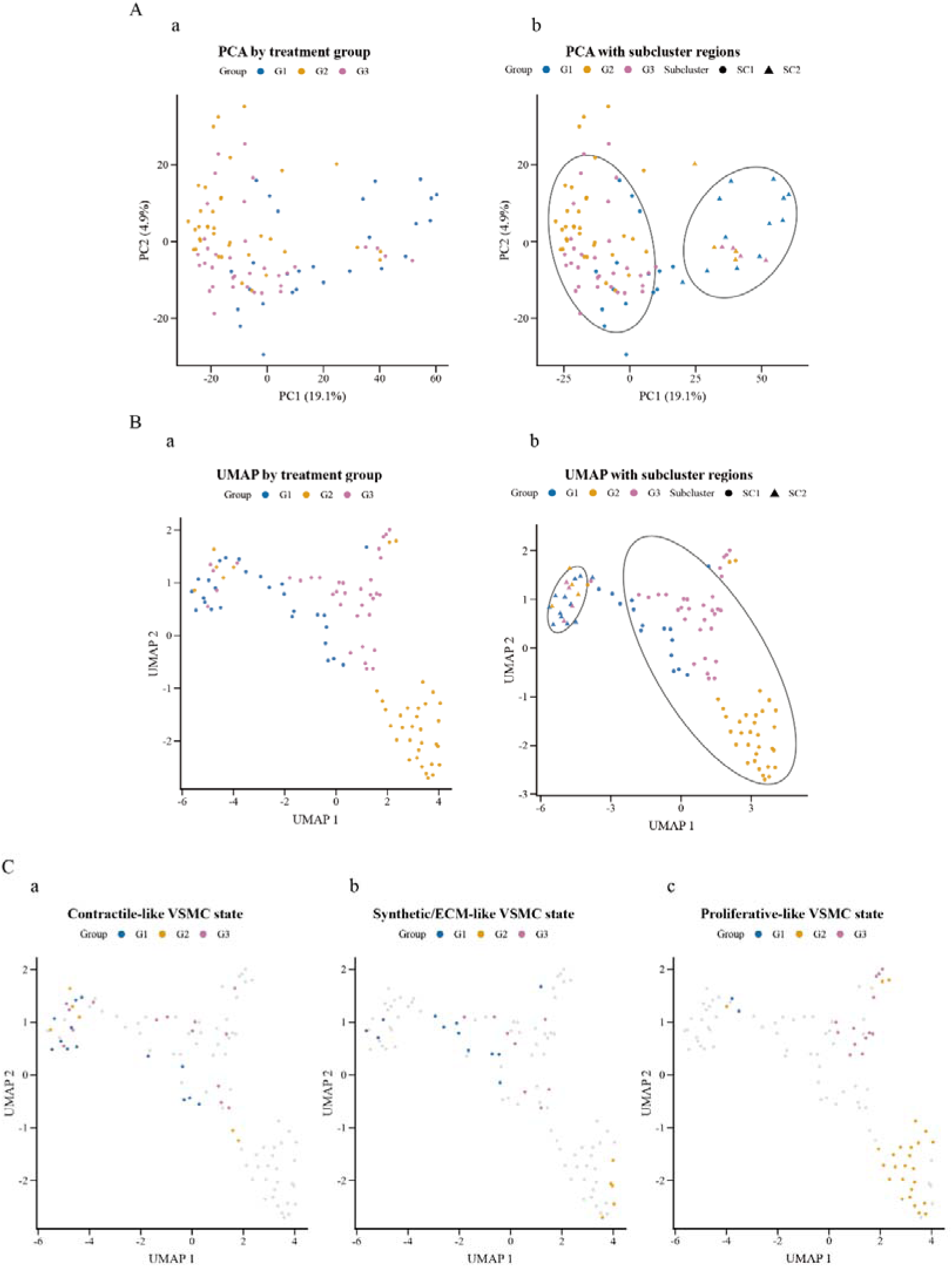
Low-dimensional VSMC state structure and functional modules. **A**, PCA coloured by treatment (**a**) and annotated with SC1/SC2 state shapes and contours (**b**). **B**, Corresponding UMAP views. **C**, Cells with high contractile-like (**a**), synthetic/ECM-like (**b**) and proliferative-like (**c**) module scores. Treatment groups are not enclosed by contours.

**Supplementary Figure S10:**
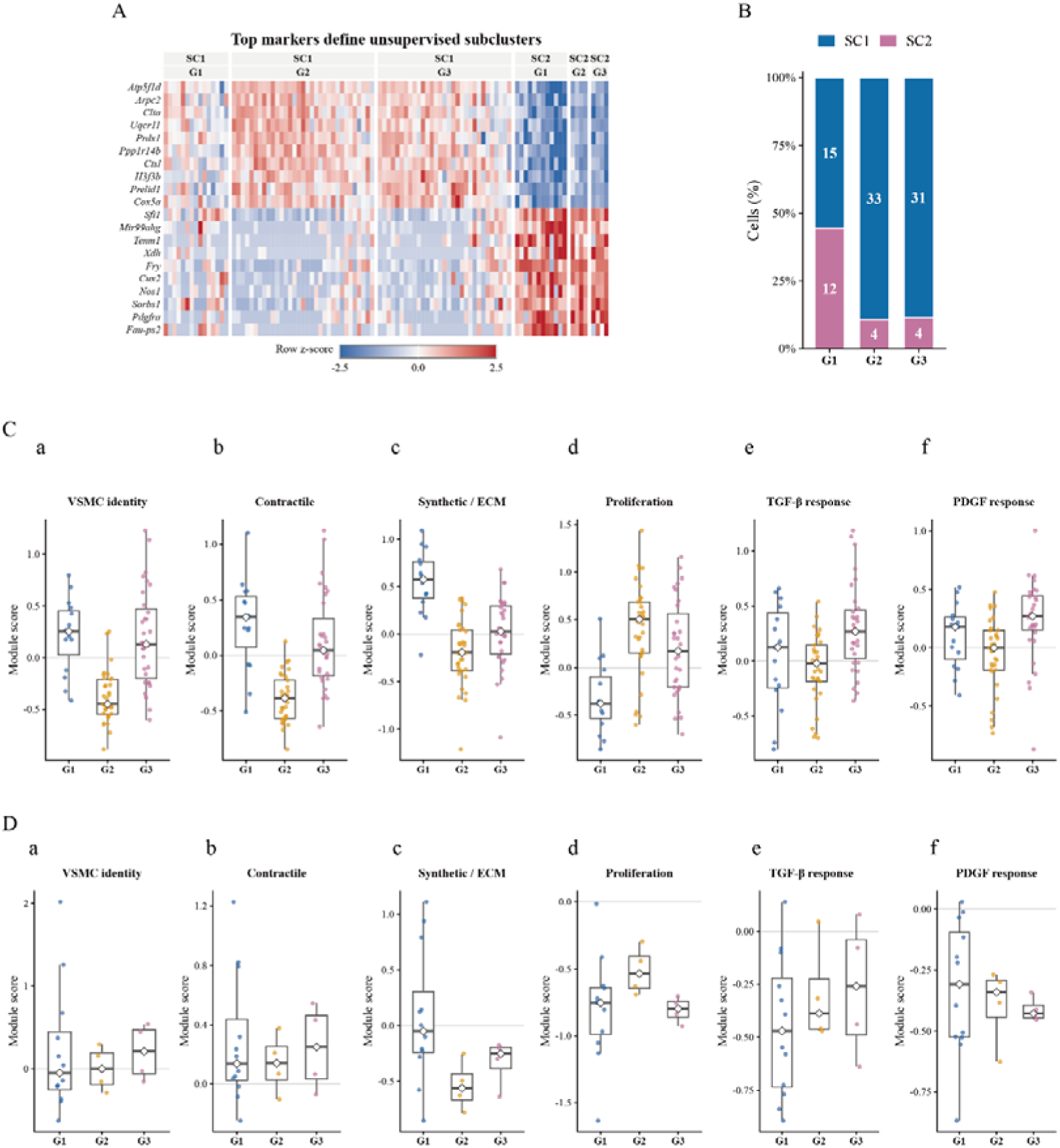
Markers, composition and within-state programmes of SC1 and SC2. **A**, Representative SC1/SC2 marker heat map. **B**, SC1/SC2 composition in G1, G2 and G3. **C,D**, VSMC identity, contractile, synthetic/ECM, proliferation, TGF-β-response and PDGF-response module scores within SC1 (**C**) and SC2 (**D**). G2 and G3 each contain four SC2 cells; these panels are descriptive.

**Supplementary Figure S11:**
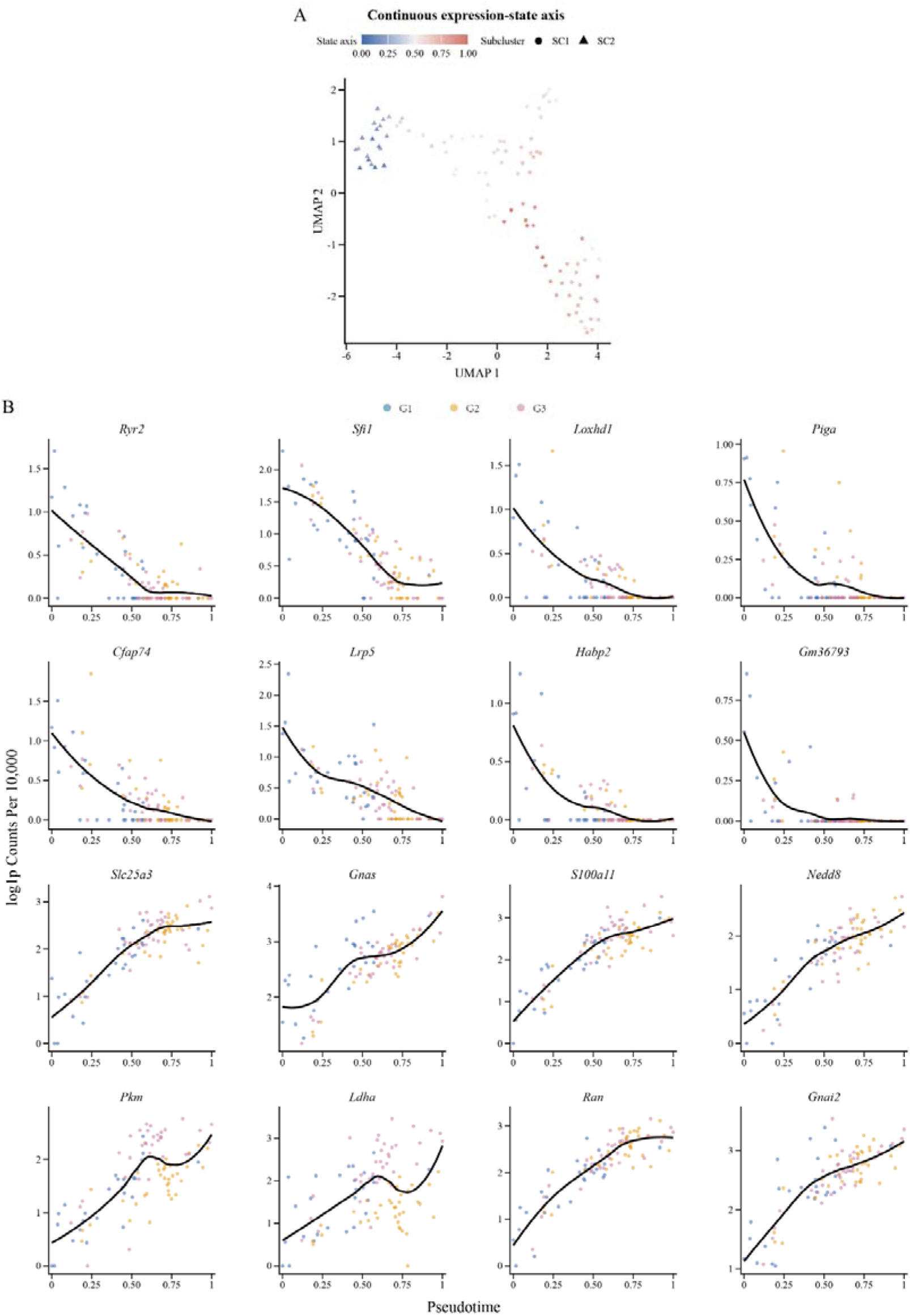
The unadjusted continuous VSMC expression-state axis. **A**, Unadjusted state-axis position in UMAP space. **B**, Sixteen representative genes varying along the axis. The axis is strongly correlated with ribosomal fraction and is not interpreted as experimental time or lineage.

**Supplementary Figure S12:**
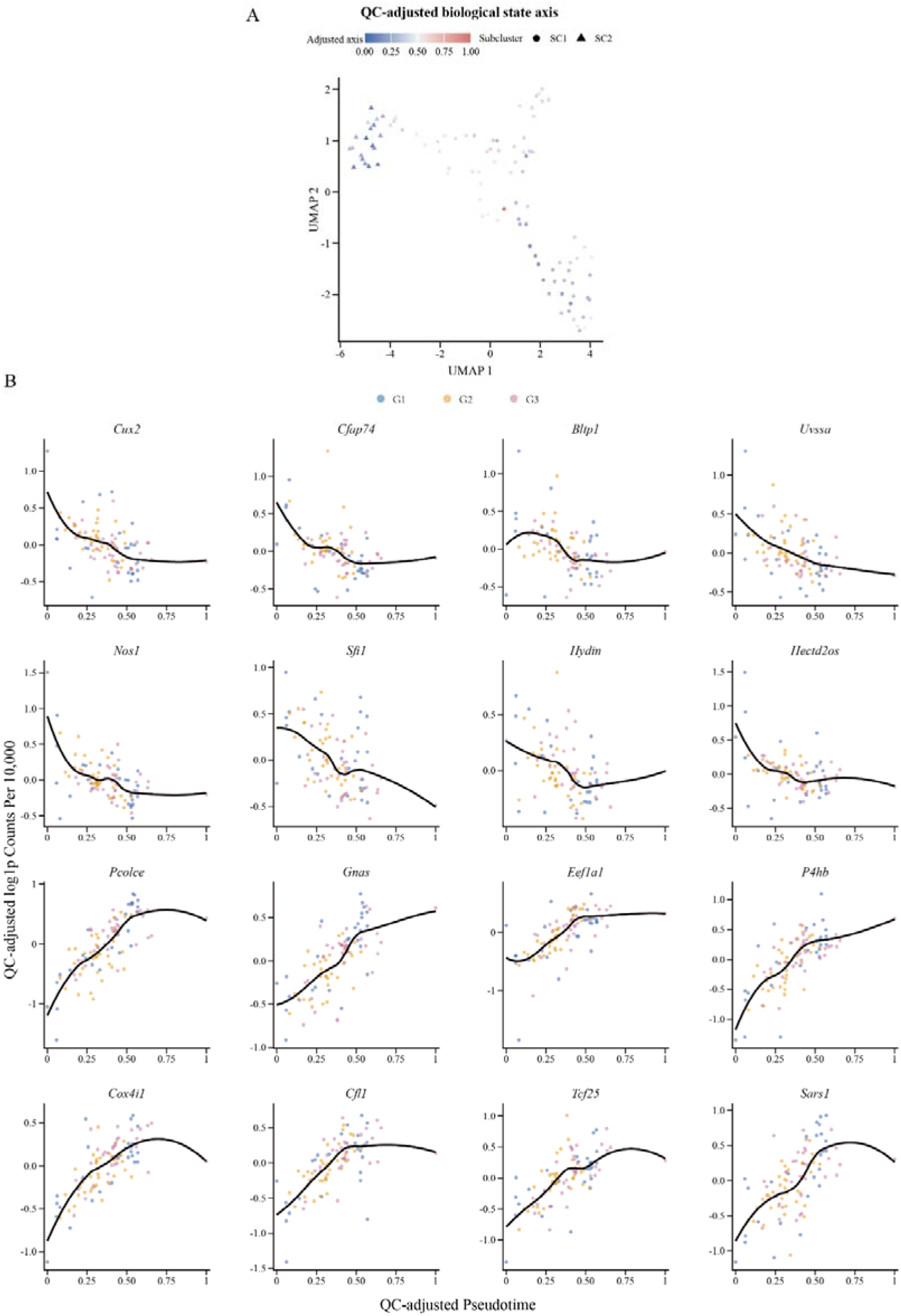
A quality-adjusted continuous VSMC expression-state axis. **A**, State axis after adjustment for UMI depth, detected genes, mitochondrial fraction, ribosomal fraction and technical library. **B**, Sixteen representative dynamic genes. Curves show LOESS trends and do not imply temporal or causal ordering.

**Supplementary Figure S13:**
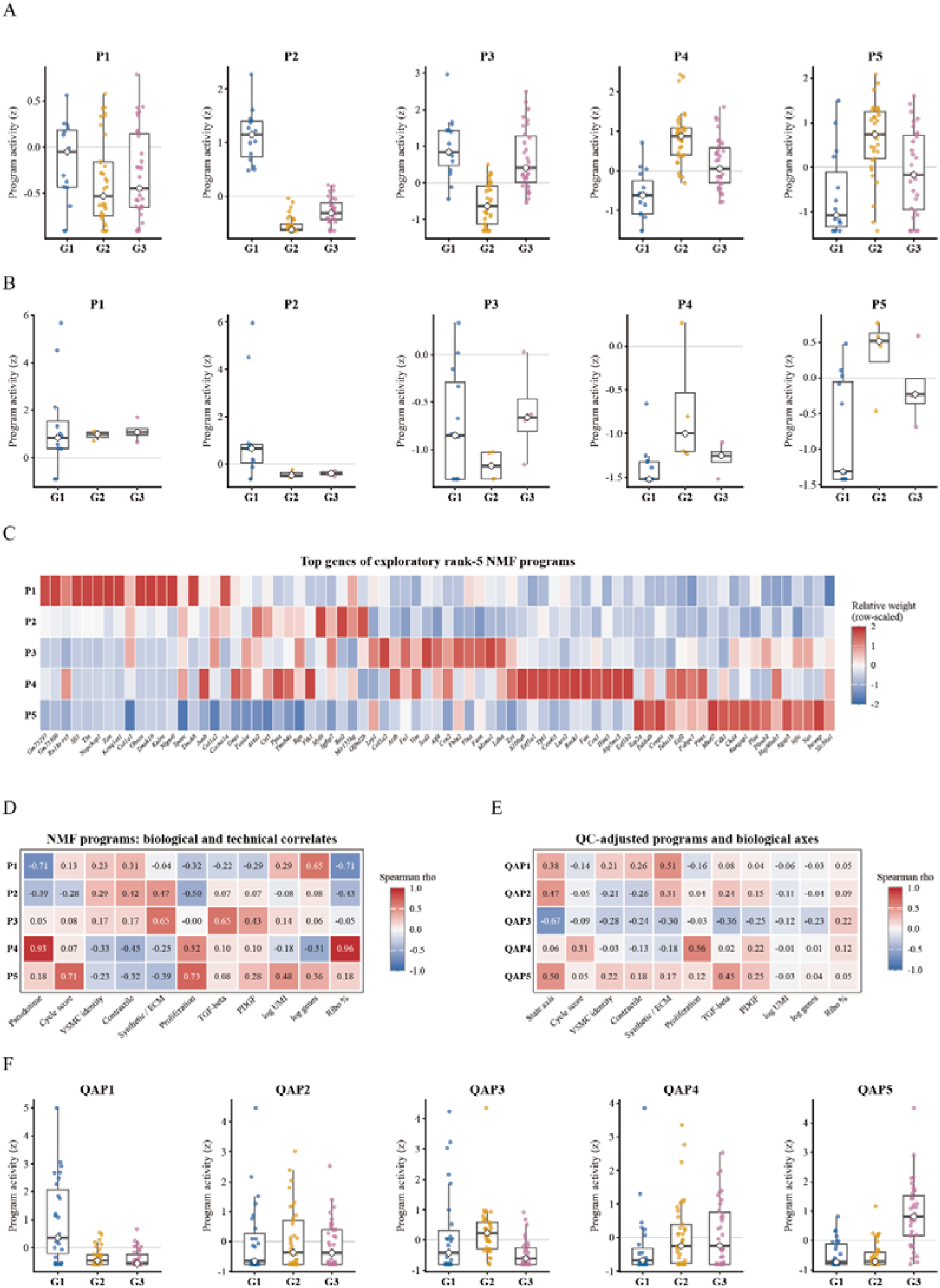
**Raw NMF and quality-adjusted expression programmes. A,B**, Activities of raw rank-5 programmes P1–P5 within SC1 (**A**) and SC2 (**B**). **C**, High-weight genes in the exploratory rank-5 model. **D**, Correlations of raw programmes with state, cell cycle, functional modules and quality metrics. **E**, Corresponding correlations for QAP1–QAP5. **F**, QAP activities across G1, G2 and G3. Programmes may coexist within a cell and are not cell types.

**Supplementary Figure S14:**
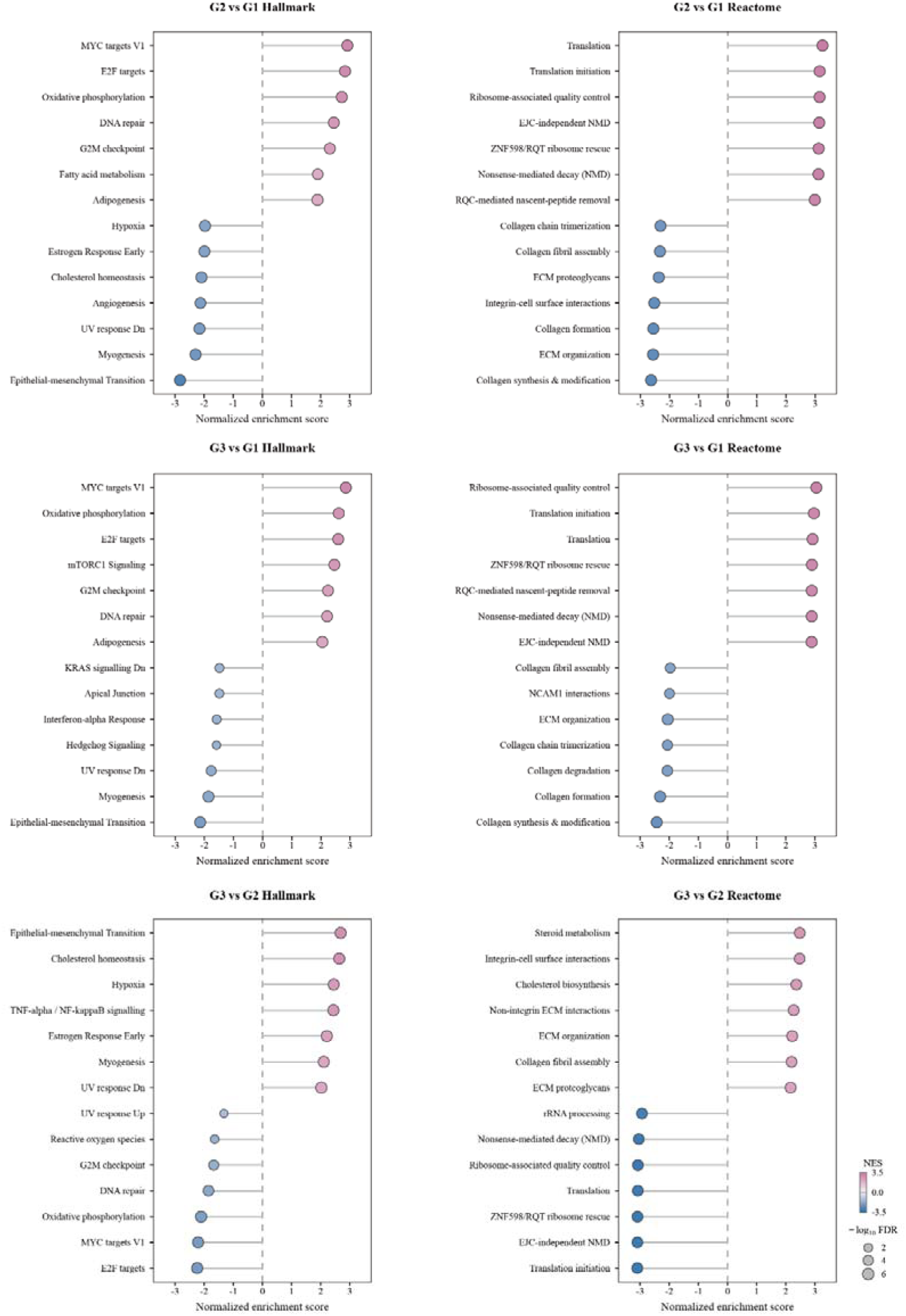
Hallmark and Reactome enrichment across treatment contrasts. Preranked enrichment results for G2 versus G1, G3 versus G1 and G3 versus G2. Positive NES denotes enrichment in the first group of each contrast.

**Supplementary Figure S15:**
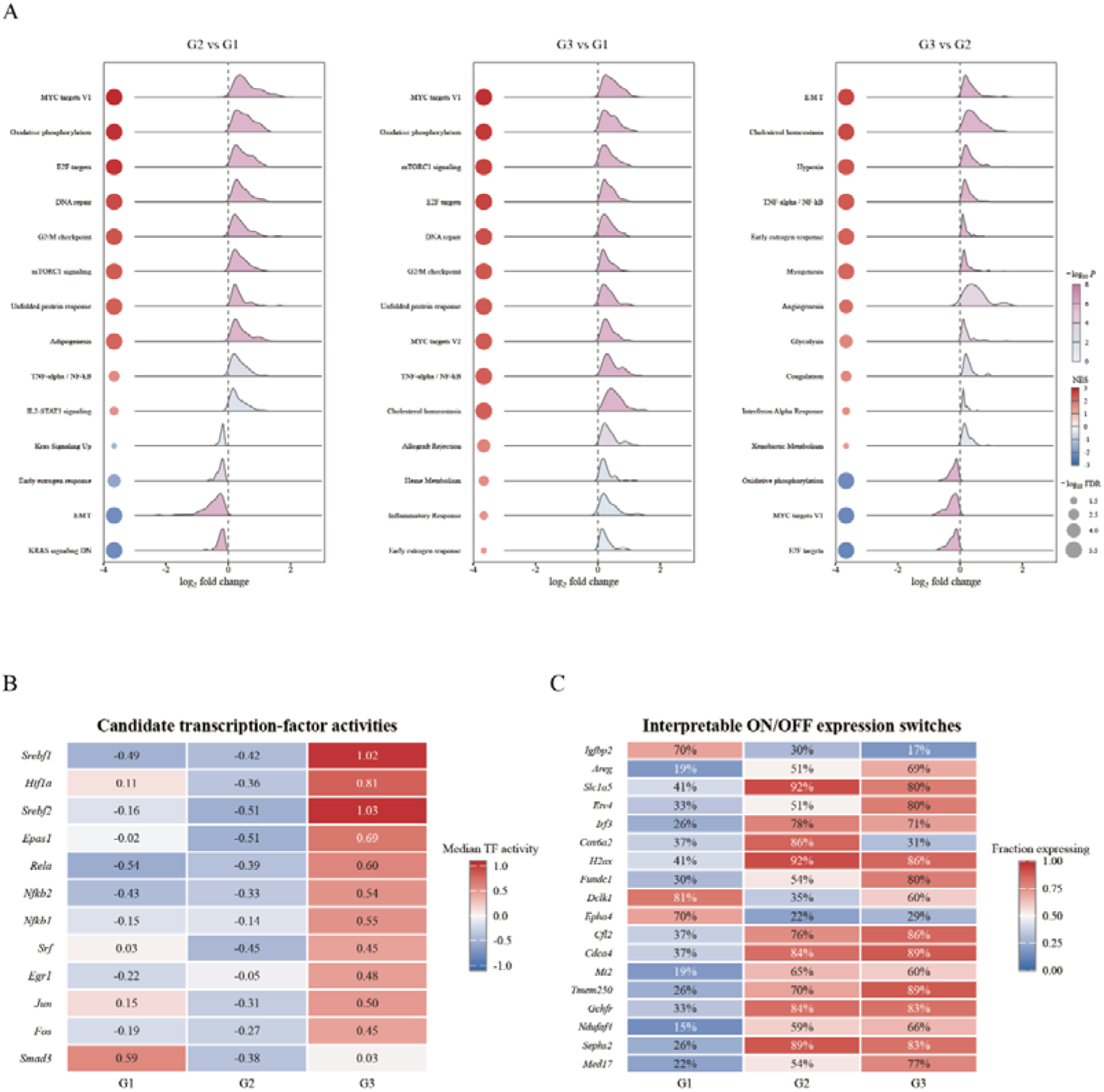
GSEA context, inferred regulators and expressing-cell fractions. **A**, Hallmark GSEA ridge-bubble plots for all three treatment contrasts on common axes. **B**, Median standardised activity of candidate transcription factors in G1, G2 and G3. **C**, Expressing-cell fractions for representative response genes.

**Supplementary Figure S16:**
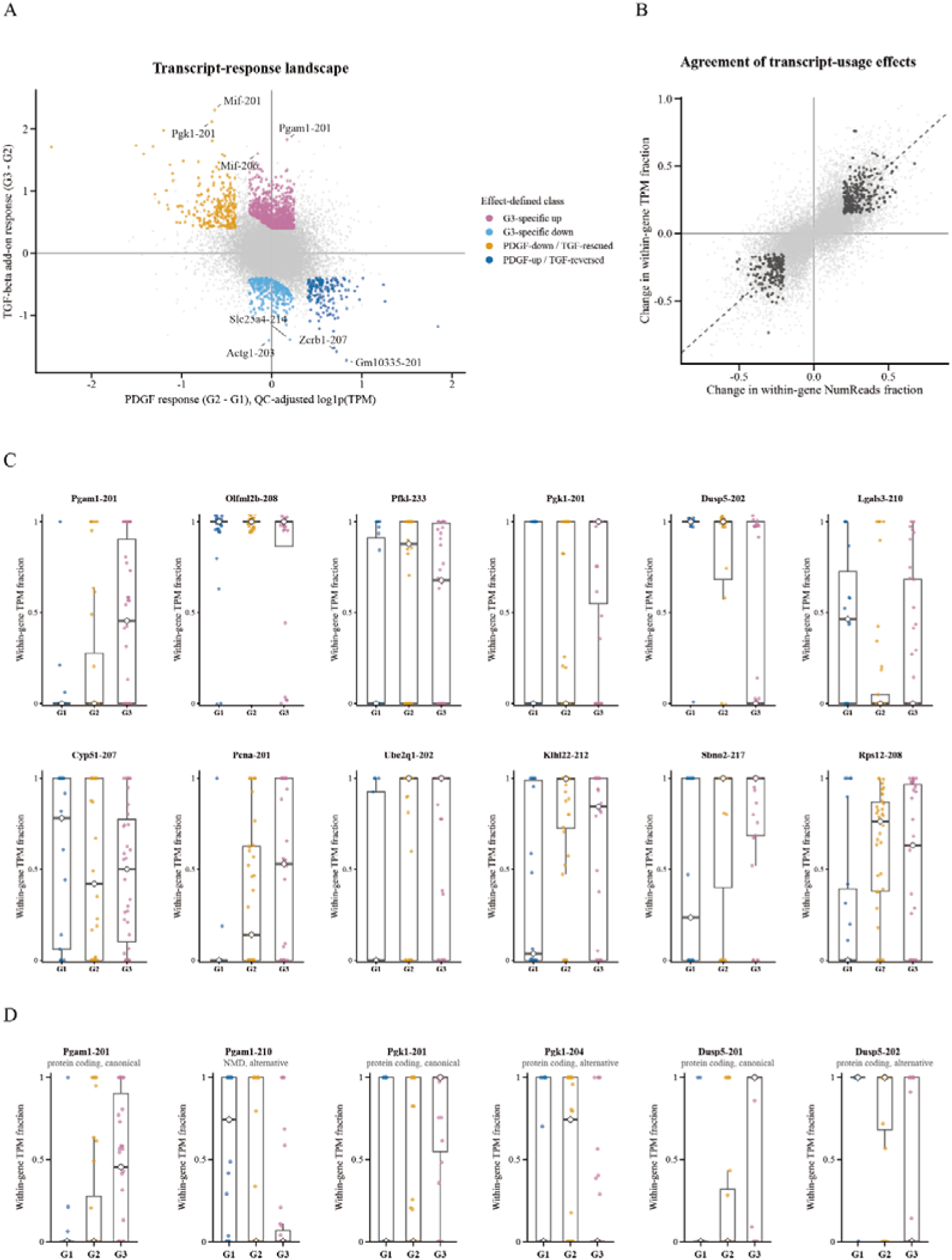
Transcript-response structure and prioritised isoform-usage candidates. **A**, Quality-adjusted transcript-response map. **B**, Concordance of within-gene usage effects derived from NumReads and TPM. **C**, Single-cell TPM usage for 12 prioritised transcript candidates. **D**, Coordinated usage of Pgam1-201/Pgam1-210, Pgk1-201/Pgk1-204 and Dusp5-201/Dusp5-202, with transcript structures for assay design.

